# PADAPT 1.0 – the Pannonian Database of Plant Traits

**DOI:** 10.1101/2022.12.05.519136

**Authors:** Judit Sonkoly, Edina Tóth, Nóra Balogh, Lajos Balogh, Dénes Bartha, Kinga Bata, Zoltán Bátori, Nóra Békefi, Zoltán Botta-Dukát, János Bölöni, Anikó Csecserits, János Csiky, Péter Csontos, István Dancza, Balázs Deák, Zoltán Konstantin Dobolyi, Anna E-Vojtkó, Ferenc Gyulai, Alida Anna Hábenczyus, Tamás Henn, Ferenc Horváth, Mária Höhn, Gusztáv Jakab, András Kelemen, Gergely Király, Szabolcs Kis, Gergely Kovacsics-Vári, András Kun, Éva Lehoczky, Attila Lengyel, Barbara Lhotsky, Viktor Löki, Balázs András Lukács, Gábor Matus, Andrea McIntosh-Buday, Attila Mesterházy, Tamás Miglécz, Attila Molnár V., Zsolt Molnár, Tamás Morschhauser, László Papp, Patrícia Pósa, Tamás Rédei, Dávid Schmidt, Ferenc Szmorad, Attila Takács, Júlia Tamás, Viktor Tiborcz, Csaba Tölgyesi, Katalin Tóth, Béla Tóthmérész, Orsolya Valkó, Viktor Virók, Tamás Wirth, Péter Török

## Abstract

We present PADAPT 1.0, the Pannonian Database of Plant Traits which relies on regional data sources and integrates existing data and new measurements on a wide range of traits and attributes of the plant species of the Pannonian Biogeographical Region and makes it freely accessible at www.padapt.eu. The current version covers the species of the region occurring in Hungary (cc. 90% of the region’s flora) and consists of 126,337 records on 2745 taxa. There are 53 plant attributes in PADAPT 1.0 organised in six major groups: (i) Habitus and strategy, (ii) Reproduction, (iii) Kariology, (iv) Distribution and conservation, (v) Ecological indicator values, and (vi) Leaf traits. By including species of the eastern part of Europe not covered by other databases, PADAPT can facilitate studying the flora and vegetation of the eastern part of the continent. Data collection will continue in the future and the PADAPT team welcomes researchers interested in contributing with data. The main task before an updated version of the database is to include species of the Pannonian region not covered by the current version. In conclusion, although data coverage is far from complete, PADAPT meets the longstanding need for a regional database of the Pannonian flora.

## Introduction

The trait-based approach has been significantly advancing our ecological and evolutionary understanding in various fields of research including vegetation science (e.g., Lavorel & Garnier, 2002, Török et al., 2020). To provide trait-based analyses with suitable data, several international plant trait databases have been established in the last decades. Some of them compile data for a wide range of traits at the regional scale, e.g., BiolFlor for the German flora (Klotz et al., 2002), LEDA for Northwest Europe (Kleyer et al., 2008), BROT for the flora of the Mediterranean (Tavşanoğlu & Pausas, 2018), or the BryForTrait database for the bryophytes of Central Europe (Bernhardt-Römermann et al., 2018). Other databases provide data for a specific group of traits at the global scale, e.g., SID, the Seed Information Database (Royal Botanical Gardens Kew, 2008), D^3^, the Dispersal and Diaspore Database (Hintze et al., 2013), and SylvanSeeds, a germination database of deciduous forests (Fernández-Pascual, 2021). There are also databases covering not only the traits and attributes of a flora of a region but also its vegetation, such as PLADIAS, the Database of the Czech Flora and Vegetation (Chytrý et al., 2021). The most frequently used (meta-)database, TRY, incorporates several databases, thus providing a global coverage for numerous plant traits (Kattge et al., 2020).

Although data on a broad range of traits can be relatively easily retrieved from these databases, it is a shortcoming that they incorporate records from several different sources and from regions with markedly different climatic conditions, often with varying measurement standards. Climate and local abiotic conditions can cause substantial intraspecific trait variability (Albert et al., 2010), which renders the application of such broad-scale databases problematic for regional-scale studies. Moreover, the existing European databases’ geographical coverage is limited and mostly focused either on the flora of the western and northwestern part of Europe, or cover the southern, mostly Mediterranean parts of the continent (Cerabolini et al., 2010, Tavşanoğlu & Pausas, 2018).

The Pannonian Biogeographical Region is situated in the eastern part of Central Europe surrounded by the Carpathians, the Alps, and the Dinaric Mountains. The whole territory of Hungary is included in this region, along with some lowland areas of the Czech Republic, Slovakia, Ukraine, Romania, and Serbia (EEA 2016, Fig. 1). Due to the above-mentioned geographical focus of the existing databases, a great proportion of the Pannonian flora is not represented in these. These problems limit the applicability of the existing databases for not just studies of the Pannonian flora and vegetation, but for studies in the eastern and central part of the continent in general, and also for studies attempting large-scale comparisons across regions.

**Figure 1.**
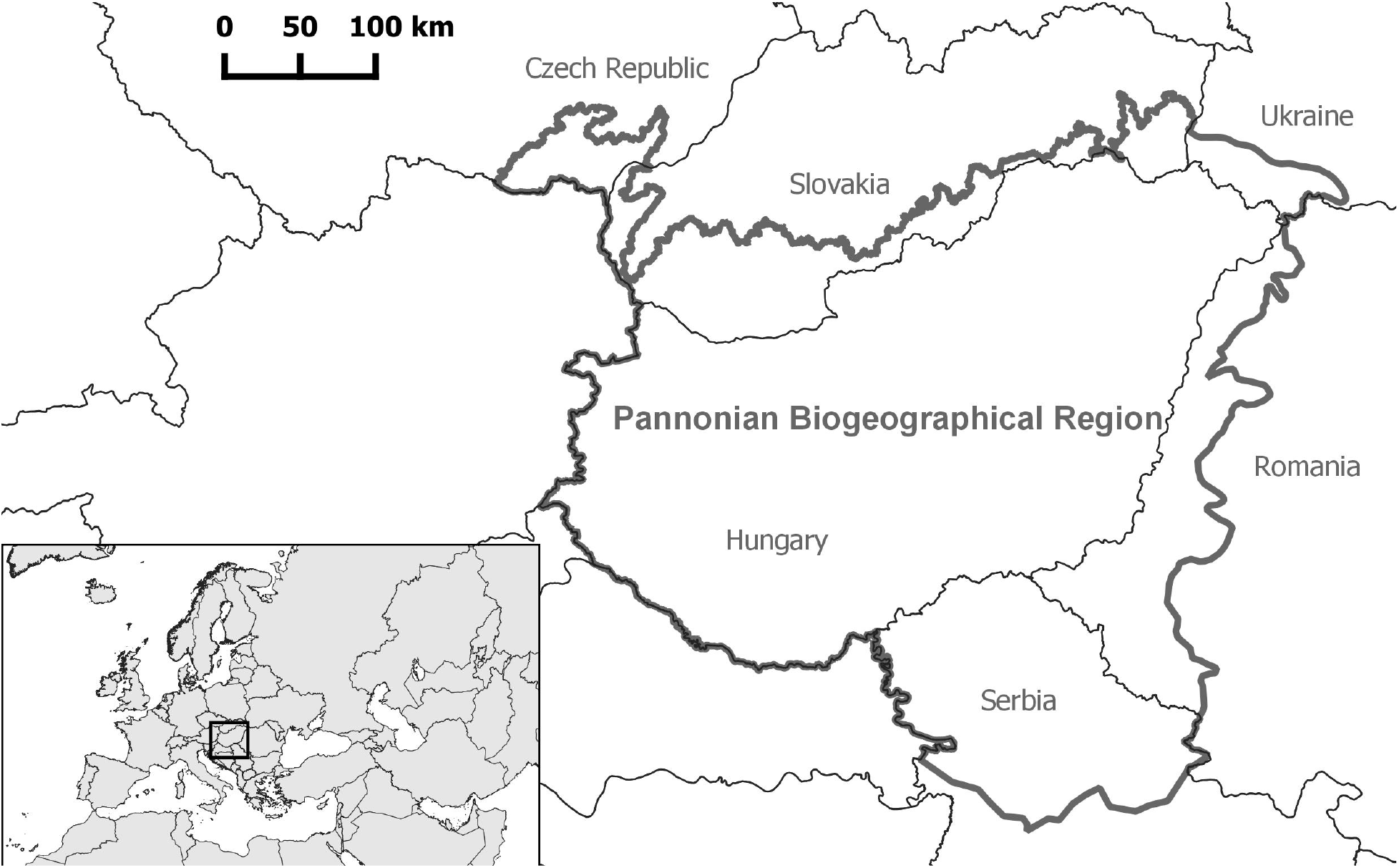
Map showing the position of the Pannonian Biogeographical Region in Europe as established by the European Environment Agency (EEA 2016).

The above-mentioned issues have motivated the compilation of a database focusing on the Pannonian flora. Here, we introduce PADAPT 1.0, the Pannonian Database of Plant Traits, which relies on regional data sources and integrates existing data and new measurements on a wide range of traits and attributes of the Pannonian flora and makes it broadly available to the international scientific community. To test whether regionally measured trait data from the Pannonian Biogeographical Region differ from trait data of the same species originating from the western part of Europe, we compare functional trait data from PADAPT 1.0 with data from the LEDA Traitbase (Kleyer et al., 2008).

### Compilation of the database

The project was initialised in 2019 during a workshop held in Debrecen, Hungary, where several Hungarian ecologists and botanists expressed their support for the creation of the database. Along with the initialisation of collecting existing data, the PADAPT team started a sampling campaign in the Pannonian region to allow for additional measurements of leaf traits and seed mass. This has resulted in new leaf trait records for 1156 species (McIntosh-Buday et al., 2022) and new thousand-seed mass (TSM) data for 114 species and counting. The PADAPT protocol for measuring seed mass and leaf traits was based on data standards for LEDA (Kleyer et al., 2008) and the protocols by Perez-Harguindeguy et al. (2016).

The current version of the database, PADAPT 1.0 only includes species of the Pannonian region that occur also in Hungary. We plan to expand the database with those Pannonian species, which occur only outside of the territory of Hungary in the future, even though there is only a small number of such species. Although the number of species in the Pannonian flora is not established in the literature, we can reasonably assume that 90% of the flora of the region is already represented in the current version. The checklist of taxa was largely based on the checklist of the Distribution atlas of vascular plants of Hungary (Bartha et al., 2021), with slight modifications to account for changes in taxonomy or nomenclature, or to exclude extinct or irrelevant taxa. Subspecies are generally not distinguished in the database, except in cases when just one subspecies occurs in Hungary (e.g., *Astragalus vesicarius* subsp. *albidus*).

A trait or attribute was included in PADAPT 1.0 if data from regional sources were available for it for a considerable share of the species and if it was considered meaningful for future ecological studies.

### Description of PADAPT 1.0

The current version, PADAPT 1.0 consists of 126,337 individual records on 2745 taxa based on 116 different published sources (for data sources see Appendix S1–S6). Most of the data are directly taken from these published sources (e.g., ecological indicator values, leaf traits or thousand-seed mass). The database also contains records specifically prepared for PADAPT based on literature sources and expert knowledge (e.g., modified Ellenberg–Müller-Dombois life form system or the phytosociological categorisation for some of the previously not covered species). A substantial part of the leaf trait and thousand-seed mass records is based on measurements carried out by the PADAPT team since the launching of the database project.

There are 53 attributes included in PADAPT 1.0, which are organised in the following six major groups: (i) Habitus and strategy, (ii) Reproduction, (iii) Kariology, (iv) Distribution and conservation, (v) Ecological indicator values, and (vi) Leaf traits. A summary of all the included traits and attributes is presented in Table 1, while detailed descriptions and data sources are available in Appendix S1–S6.

**Table 1.** Overview of the 53 attributes included in PADAPT 1.0 organised in six major groups.

|  | Attribute | Type | Units | Range | Data coverage |
| --- | --- | --- | --- | --- | --- |
| Habitus and Strategy | Minimum height | Cont. | cm | 1–3000 | 95.6% |
|  | Maximum height | Cont. | cm | 3–4500 | 95.9% |
|  | Life form (Raunkiær) | Categ. | – | * | 96.2% |
|  | Social Behaviour Type (Borhidi) | Categ. | – | * | 80.7% |
|  | Ujvárosi's life form system | Categ. | – | * | 28.8% |
|  | Modified Ellenberg–Müller-Dombois life form system | Categ. | – | * | 89.6% |
| Reproduction | Onset of flowering | Ord. | Months | 1–11 | 96.1% |
|  | Duration of flowering | Ord. | Months | 1–12 | 96.1% |
|  | Thousand-seed mass (TSM) average | Cont. | g | 0.001–5002.4 | 67.1% |
|  | Seed mass category (Csontos) | Ord. | – | 1–8 | 80.7% |
|  | Seed bank persistence index | Cont. | – | 0–1 | 19.7% |
|  | Dispersal strategy (Sádlo et al.) | Categ. | – | * | 100% |
| Kariology | Chromosome number | List** | – | ** | 83.7% |
|  | Ploidy level | Categ. | – | * | 3.1% |
| Distribution and Conservation | Phytosociological categorization (Soó) | Categ. | – | * | 73.2% |
|  | Phytosociological categorization (Borhidi) | Categ. | – | * | 75.0% |
|  | Nativeness | Categ. | – | * | 100% |
|  | Residence time | Categ. | – | * | 100% |
|  | Introduction mode | Categ. | – | * | 100% |
|  | Invasion status | Categ. | – | * | 100% |
|  | Area type | Categ. | – | * | 82.2% |
|  | Conservation status | Categ. | – | * | 100% |
|  | Conservation value | Categ. | HUF | * | 100% |
|  | Year of protection | Categ. | Year | * | 100% |
|  | IUCN Red List category | Categ. | – | * | 100% |
|  | Hungarian Red List category | Categ. | – | * | 100% |
|  | Conservation value category (Simon) | Categ. | – | * | 99.7% |
| Ecological indicator values | Ellenberg's L | Ord. | – | 1–9 (and ×) | 64.5% |
|  | Ellenberg's T | Ord. | – | 1–9 (and ×) | 64.5% |
|  | Ellenberg's K | Ord. | – | 1–9 (and ×) | 64.5% |
|  | Ellenberg's F | Ord. | – | 1–9 (and ×) | 64.5% |
|  | Ellenberg's R | Ord. | – | 1–9 (and ×) | 64.5% |
|  | Ellenberg's N | Ord. | – | 1–9 (and ×) | 64.5% |
|  | Ellenberg's S | Ord. | – | 0–9 | 64.5% |
|  | Zólyomi's T | Ord. | – | 1–7 (and 0) | 62.5% |
| Ecological indicator values<br>(cont.) | Zólyomi's W | Ord. | – | 0–11 | 63.4% |
|  | Zólyomi's R | Ord. | – | 1–5 (and 0) | 62.2% |
|  | Soó's T | Ord. | – | 1–5 (and 0) | 76.9% |
|  | Soó's F | Ord. | – | 1–5 (and 0) | 77.0% |
|  | Soó's R | Ord. | – | 1–5 (and 0) | 75.0% |
|  | Soó's N | Ord. | – | 1–5 (and 0) | 72.8% |
|  | Borhidi's T | Ord. | – | 0–8 | 77.1% |
|  | Borhidi's W | Ord. | – | 1–12 | 77.1% |
|  | Borhidi's R | Ord. | – | 1–9 | 77.1% |
|  | Borhidi's N | Ord. | – | 1–9 | 77.1% |
|  | Borhidi's L | Ord. | – | 1–9 | 77.1% |
|  | Borhidi's C | Ord. | – | 1–9 | 77.1% |
|  | Borhidi's S | Ord. | – | 0–9 | 77.1% |
| Leaf traits | Leaf area (LA) | Cont. | mm<sup>2</sup> | 2.001–<br>194,933.617 | 57.8% |
|  | Leaf dry matter content (LDMC) | Cont. | mg/g | 4.049–<br>636.872 | 57.4% |
|  | Specific leaf area (SLA) | Cont. | mm<sup>2</sup>/mg | 3.602–356.0 | 57.4% |
|  | Leaf fresh mass | Cont. | g | 0.001–84.294 | 55.6% |
|  | Leaf dry mass | Cont. | mg | 0.1–9743.3 | 55.4% |

Regarding the species included in the current version, data coverage is complete (100%) for dispersal strategy, nativeness, residence time, introduction mode, invasion status, conservation status, conservation value (HUF), year of protection, IUCN Red List category, and Hungarian Red List Category. Data coverage is at least 80% for about 40% of the attributes and at least 50% for almost all attributes (>94% of attributes) (Table 1).

As data for some traits (e.g., thousand-seed mass) were gathered from different literature sources, species can have multiple records. In these cases, trait data can be downloaded either as all data from a single source or as the average value for each species from different sources. A derivate value can be obtained in the case of seed bank persistence index that is calculated based on seed bank type data from all sources.

### Comparing PADAPT with other databases

To quantify how many additional species are covered by PADAPT regarding functional traits compared to TRY, the biggest plant trait database (Kattge et al., 2020), we obtained datasets of (i) seed dry mass (corresponding to thousand-seed mass, TSM), (ii) leaf dry matter content (LDMC), (iii) leaf dry mass, (iv) leaf area (LA), and (v) specific leaf area (SLA) from TRY on 6^th^ September 2022. Compared to TRY, PADAPT 1.0 provides TSM data for 439 additional species, LA for 890 additional species, LDMC for 455 additional species, SLA for 529 additional species, and leaf dry mass data for 531 additional species.

We also aimed to test whether regionally measured trait data from the Pannonian region differ from trait data for the same species originating from the western part of Europe. To this end, we compared functional trait data from PADAPT 1.0 with data from the LEDA Traitbase (Kleyer et al., 2008) for thousand-seed mass (TSM), leaf area (LA), leaf dry matter content (LDMC), specific leaf area (SLA), and leaf dry mass. We decided not to compare our data with data from TRY as a considerable part of the leaf trait records included in PADAPT (those published by E-Vojtkó et al. 2020) are also included in TRY.

Data files for the following traits were obtained from LEDA on 23^rd^ November 2021: (i) Seed mass (corresponding to TSM in PADAPT 1.0), (ii) Leaf size (corresponding to LA), (iii) SLA, (iv) LDMC, and (v) Leaf mass (corresponding to leaf dry mass after converting mg to g). We extracted data from LEDA data files for the species also present in our database and calculated the mean value of all records from LEDA for each species. To maximise comparability, we considered only those records that were described as ‘actual measurement’. In the case of leaf traits, we excluded measurements without petiole and rachis, and in the case of seed mass, we excluded measurements of multi-seeded generative dispersules. We then compared the data using pairwise Wilcoxon signed-rank tests in R version 4.0.3 (R Core Team 2020).

TSM, SLA and LDMC values in PADAPT significantly differed from those in LEDA (Table 2, Fig. 2A, 2C and 2D, respectively), while LA and leaf dry mass values did not differ significantly (Table 2, Fig. 2B and 2E, respectively). The considerably different TSM, SLA and LDMC values in PADAPT compared to LEDA may be attributed to the drier, more continental climate of the Pannonian region compared to the western part of Europe, as plants inhabiting regions with drier climates tend to have thicker leaves with low SLA and high LDMC values (e.g., Fonseca et al., 2000; Wright et al., 2004). Although some results suggest that there is a negative relationship between precipitation and seed mass (Baker, 1972; Wright & Westoby, 1999), more recent studies found that seed mass decreases with increasing aridity and precipitation variability (Harel et al. 2011, Yu et al. 2007).

**Table 2.** Comparison of data from the LEDA Traitbase and PADAPT 1.0 (pairwise Wilcoxon signed-rank tests).

|  | LEDA<br>(mean±SE) | PADAPT<br>(mean±SE) | *p* value | *n* |
| --- | --- | --- | --- | --- |
| **Thousand-seed mass (TSM)** | 18.79±5.06 | 16.19±4.60 | 6.14e-12 | 1246 |
| **Leaf area (LA)** | 2597.78±261.24 | 2231.02± 220.74 | 0.1012 | 495 |
| **Specific leaf area (SLA)** | 24.92±0.39 | 24.48± 0.47 | 5.862e-05 | 960 |
| **Leaf dry matter content (LDMC)** | 201.59±3.75 | 251.55± 4.59 | <2.2e-16 | 476 |
| **Leaf dry mass** | 146.47± 18.94 | 132.32± 17.49 | 0.2146 | 491 |

**Figure 2.**
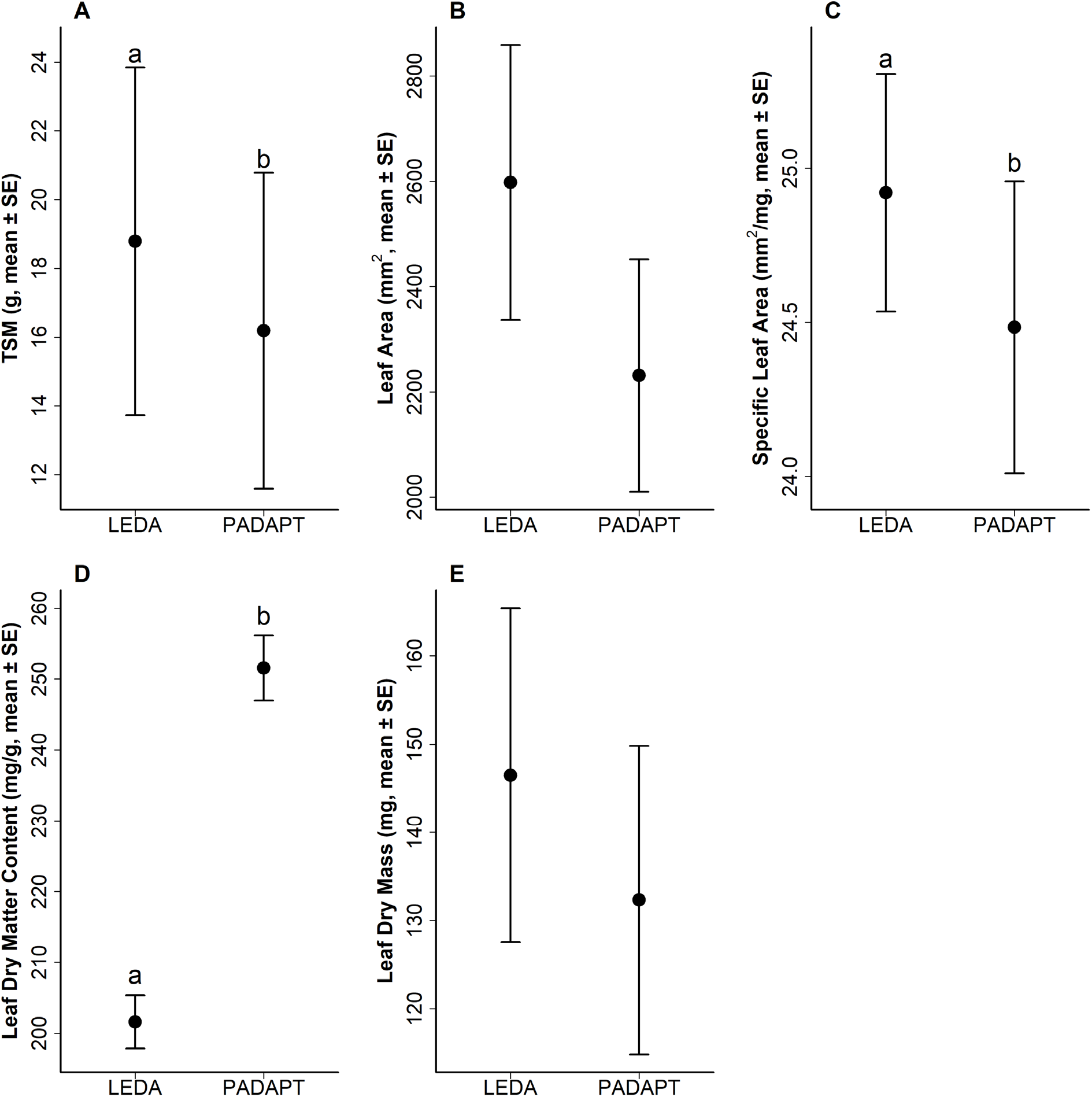
Comparison of data from the LEDA Traitbase and PADAPT 1.0 for (A) thousand-seed mass, (B) leaf area, (C) specific leaf area, (D) leaf dry matter content and (E) leaf dry mass. Wilcoxon signed-rank tests, significant differences are denoted by different letters.

### Data access and usage

The database is freely accessible online at www.padapt.eu. The PADAPT website has two language options: English and Hungarian. Under the Species menu, one can access species’ pages where all data describing a species are shown along with photo(s) and a map showing its distribution in Hungary (Fig. 3). We plan to prepare distribution maps for the whole Pannonian Biogeographical Region for an updated version of the database. To access a species’ page, one can use the search bar or click on a family from the list which retrieves a list of genera, then clicking on a genus retrieves all available species belonging to that genus.

**Figure 3.**
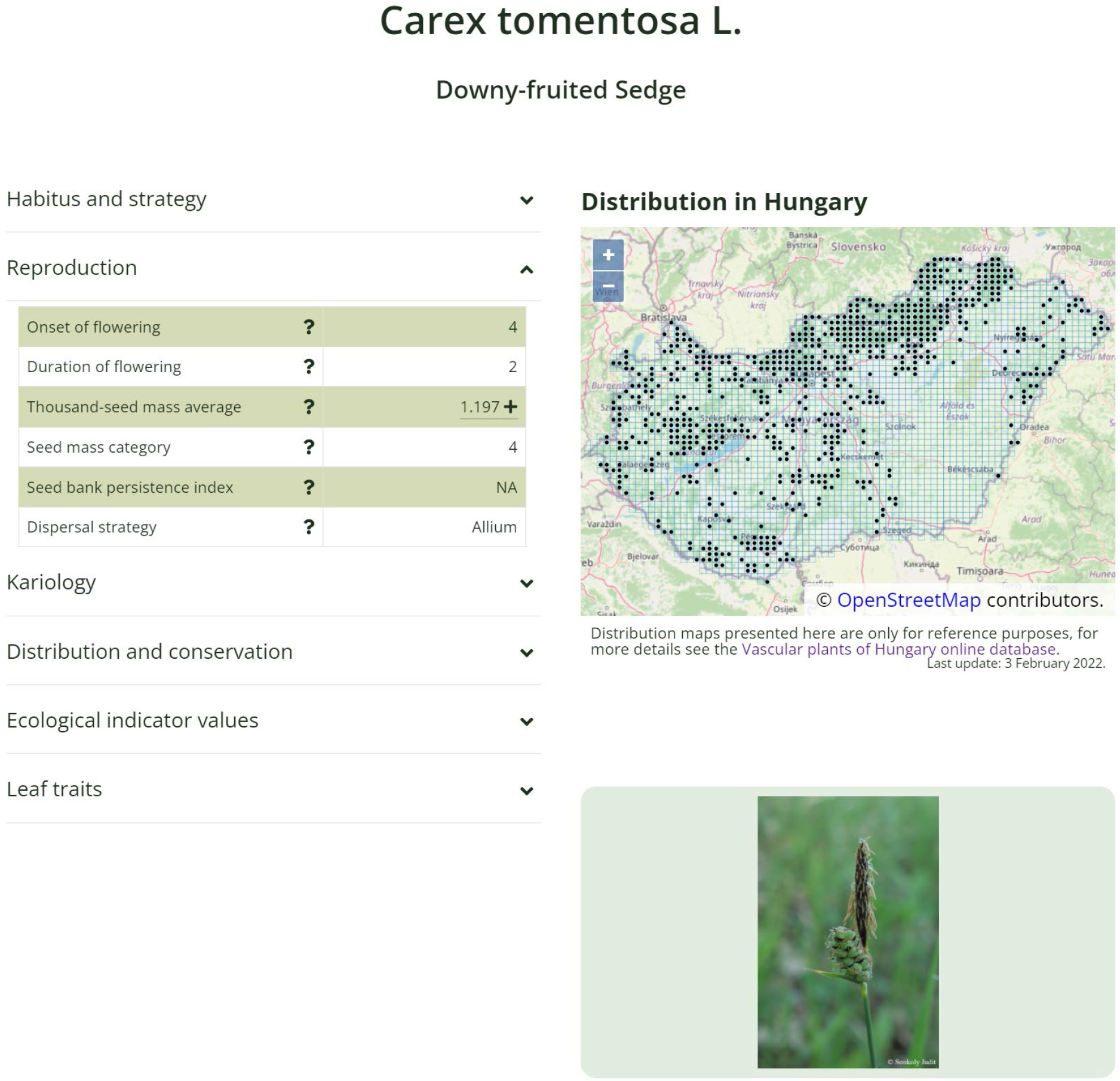
Example for a species’ page at www.padapt.eu where all data describing a species is available along with photo(s) and a map showing its distribution in Hungary.

There are two options to download data from the website. Under the Download menu one can either choose an attribute and download all data for that attribute, or search for a species and download all data for that species. Data can be downloaded in the form of separate CSV files.

All data included in PADAPT 1.0 are public, but the present paper along with any source publication provided should be appropriately referenced when using the data. Information on what and how to cite when using data for an attribute from PADAPT can be found on the attribute information pages under the Info menu. Attribute information pages also provide descriptions for all attributes. Descriptions and data sources are also available in Appendix S1–S6.

## Conclusion and outlook

Although data coverage is far from complete for many adventives and other species recently discovered in the territory of Hungary, PADAPT 1.0 meets the longstanding need for a regional database of the Pannonian flora.

PADAPT 1.0 provides regionally collected data and covers several species with continental, Balkanic, and Pontic distributions, thus, it will promote studies of the flora and vegetation of not only the Pannonian region, but also the eastern part of the continent. The new database highlights data gaps, facilitates their targeted filling and promotes the exploration of intraspecific trait variability. By providing an example for a regional trait database, PADAPT may also promote the creation of other regional-scale databases. The importance of having regionally measured trait data is also confirmed by the fact that trait data retrieved from LEDA for our species differed significantly in the case of seed mass, specific leaf area and leaf dry matter content, presumably due to climatic differences. Thus, in the case of studies carried out in the Pannonian region and presumably also in the whole eastern part of Europe, it is highly advisable to use PADAPT instead of LEDA.

There are several possibilities and directions for the further development of the database. (i) In the coming years, we will continue the field samplings to provide trait data for the currently data-deficient species in the database. (ii) The most substantial development will be extending the species list to include species of the Pannonian Biogeographical Region missing from the flora of Hungary, as the current version (1.0) includes only the species that can be found in the territory of Hungary. Although the number of species in the Pannonian flora is not established in the literature, we estimated it by quantifying the number of species that are present in the Pannonian part of the Czech Republic according to Pladias (the database of the Czech Flora and Vegetation, Chytrý et al. 2021), but not in Hungary. Based on this, we can reasonably assume that approximately 90% of the flora of the region is already represented in the current version, thus, the species list will increase by cc. 10% in the future. (iii) Intraspecific trait variability could be captured by targeted sampling in several populations of each species from different subregions. (iv) Several other plant traits and attributes missing from the current version are planned to be included in the database, such as root traits, CSR strategy types calculated based on regionally measured traits (based on Hodgson et al. 1999 and/or Pierce et al. 2017), biologically active components and palatability for herbivores, presence and types of mycorrhizae, disturbance tolerance, seed dimensions (length and width) and seed hardness, leaf nitrogen and phosphorous content etc.

Data collection will continue in the future and the PADAPT team welcomes any researcher interested in contributing to PADAPT with new data. In the coming years we expect to release PADAPT 2.0 complemented with additional attributes and further species.

## Acknowledgements

We are grateful for Patricia Díaz Cando, Kata Frei, Alexandra Tomasovszky and Viktória Törő-Szijgyártó for their help in trait measurements.

## Author contributions

PT conceived the idea of the database and led the work of the consortium; all authors contributed with data; JS and ET coordinated the compilation of the data; JS coordinated the construction of the website; JS analysed the data; JS, ET and PT wrote the first draft of the manuscript and all authors contributed critically to the draft and gave final approval for publication. Lists of authors who worked on particular subsets of the database are given in the appendices containing the descriptions and data sources of the attributes.

## Conflict of interest

None of the authors have a conflict of interest to disclose.

## Data availability statement

All data in PADAPT 1.0 are freely available online at www.padapt.eu.

## List of appendices

**Appendix S1.** Descriptions and data sources of the attributes in the ‘Habitus and strategy’ group.

**Appendix S2.** Descriptions and data sources of the attributes in the ‘Reproduction’ group.

**Appendix S3.** Descriptions and data sources of the attributes in the ‘Kariology’ group.

**Appendix S4.** Descriptions and data sources of the attributes in the ‘Distribution and conservation’ group.

**Appendix S5.** Descriptions and data sources of the attributes in the ‘Ecological indicator values’ group.

**Appendix S6.** Descriptions and data sources of the attributes in the ‘Leaf traits’ group.

## Notes

**Funding information**, The authors and/or the database building project were supported by NKFIH: PD 137747 (JS), KH 130320 (ET), K 119225 (PT), K 124796 (ZB), K 137573 (PT), KKP 144068 (PT), PD 138859 (AL), PD 137828 (AT) and PD 138715 (VL). AK and ZB were supported by the Bolyai János Scholarship of the Hungarian Academy of Sciences (BO/00713/19 and BO/00298/21, respectively). AAH was supported by the New National Excellence Program of the Ministry for Innovation and Technology of Hungary (UNKP-21-3-SZTE-389).

### Competing Interest Statement

The authors have declared no competing interest.

