## Appendix S1 for "PADAPT 1.0 – the Pannonian Database of Plant Traits"

**Supporting Information to the paper Sonkoly, J. et al. Introducing PADAPT 1.0, the Pannonian Database of Plant Traits. *Journal of Vegetation Science.***

**Appendix S1. Descriptions and data sources of the attributes in the 'Habitus and strategy' group.**

### Plant height (minimum and maximum height)

The height of the aboveground shoot (including also flowers and other non-photosynthesizing organs), or the length of the shoots for vines, of the adult plant in centimeters. Minimum and maximum heights are the smallest and greatest typical values of adult plants.

Data source and citation:

Király G. (ed.) (2009) Új magyar füvészkönyv. Magyarország hajtásos növényei. Határozókulcsok. [New Hungarian Herbal. The Vascular Plants of Hungary. Identification key.] – Aggteleki Nemzeti Park Igazgatóság, Jósvafő, 616 p.

Sonkoly, J., Tóth, E., Balogh, N., Balogh, L., Bartha, D. ... Török, P. (2022) Introducing PADAPT 1.0, the Pannonian Database of Plant Traits. Journal of Vegetation Science (submitted manuscript)

### Life form

Raunkiær's life form of the species. Raunkiær's system is based on the perennating organs (seeds or buds), and in the latter case on the position of the buds relative to the soil or water surface during the unfavourable season.

Ph – Phanerophytes: woody plants with resting buds high above the soil surface

- **MM** – Mega-mesophanerophytes: trees
- **M** – Microphanerophytes: shrubs
- **N** – Nanophanerophytes: subshrubs

**Ch** – Chamaephytes: perennial plants with resting buds close to the soil surface (dwarf shrubs, cushion plants).

**H** – Hemicryptophytes: perennial plants with resting buds on the soil surface or right below the surface

**G** – Geophytes: perennial plants with resting buds in the soil, well below the surface

**HH** – Hydato-helophytes: aquatic plants with resting buds in the water or at the bottom of the water body

**TH** – Hemitherophytes: biennial plants that overwinter in the first year and die in the second year after fruit ripening

**Th** – Therophytes: annual plants that survive unfavourable seasons in the form of seeds

**E** – Epiphytes: plants living on other plants, mostly on trees

Data source and citation:

Király G. (ed.) (2009) Új magyar füvészkönyv. Magyarország hajtásos növényei. Határozókulcsok. [New Hungarian Herbal. The Vascular Plants of Hungary. Identification key.] – Aggteleki Nemzeti Park Igazgatóság, Jósvafő, 616 p.

Sonkoly, J., Tóth, E., Balogh, N., Balogh, L., Bartha, D. ... Török, P. (2022) Introducing PADAPT 1.0, the Pannonian Database of Plant Traits. Journal of Vegetation Science (submitted manuscript)

### Borhidi's social behaviour type (SBT)

Species are classified in the social behaviour type system of Borhidi according to the Flóra database (Horváth et al. 1995). The S, C, and G basic types combined with the rarity classification (r – rare, u – unique) generate an additional dimension.

I. – Natural competitors (**C**)

- **Cr** – Rare competitors
- **Cu** – Unique competitors

II. – Stress-tolerants (**ST**)

- Specialists (**S**)
  - **Sr** – Rare specialists
  - **Su** – Unique specialists
- Generalists (**G**)
  - **Gr** – Rare generalists
  - **Gu** – Unique generalists

III. – Ruderals (**R**)

- Natural Pioneers (**NP**), plant species of stressed habitats
- Plant species of anthropogenically disturbed habitats:
  - **DT** – Disturbance-tolerants
  - **W** – Weeds
- Anthropogenic alien plants:
  - **I** – Introduced aliens
  - **A** – Adventive aliens
- Competitors of secondary habitats:

- **RC** – Ruderal Competitors
- **AC** – Alien Competitors

Data source and citation:

Borhidi A. (1995) Social behaviour types, the naturalness and relative ecological indicator values of the higher plants in the Hungarian Flora. Acta Botanica Hungarica 39: 97-181.

Horváth, F., Dobolyi, K., Morschhauser, T., Lőkös, L., Karas, L., & Szerdahelyi, T. (1995) Flóra adatbázis 1.2. Taxon-lista és attribútum állomány. Vácrátót: MTA ÖBKI. [Flora database 1.2, List of taxa and attributes.]

Sonkoly, J., Tóth, E., Balogh, N., Balogh, L., Bartha, D. ... Török, P. (2022) Introducing PADAPT 1.0, the Pannonian Database of Plant Traits. Journal of Vegetation Science (submitted manuscript)

**Ujvárosi's life form system**

The life-form system of weeds for Hungarian conditions was prepared by Ujvárosi (1973).

Annuals (Th):

- $\mathbf{T_1}$: Species germinating in autumn and setting seed in spring
- $\mathbf{T_2}$: Species germinating in autumn and setting seed in early summer
- $\mathbf{T_3}$: Species germinating in spring and setting seed in early summer
- $\mathbf{T_4}$: Species germinating in spring and setting seed in autumn

**HT**: Biennials

Perennials:

Perennials with underground and aboveground overwintering stems:

- **Ch**: with a stem <30 cm
- **Ph**: with a stem >30 cm

Perennials with aboveground stem dying down:

Perennials overwintering at ground level (H)

- $\mathbf{H_1}$: Perennials overwintering at ground level, with a tufted root system
- $\mathbf{H_2}$: Perennials overwintering at ground level, propagating with stolons
- $\mathbf{H_3}$: Perennials overwintering at ground level, with taproots capable of propagation
- $\mathbf{H_4}$: Perennials overwintering at ground level, with taproots not capable of propagation
- $\mathbf{H_5}$: Perennials overwintering at ground level, with slanting rhizomes

Perennials overwintering underground (G)

- $\mathbf{G_1}$: Perennials overwintering underground with underground stolons
- $\mathbf{G_2}$: Perennials overwintering underground with tubers
- $\mathbf{G_3}$: Perennials overwintering underground with rhizomes

- **$G_4$**: Perennials overwintering underground with bulbs

**HY:** Aquatic plants (hydatophytes)

Data source and citation:

Ujvárosi, M. (1973) Gyomnövények. Mezőgazdasági Kiadó, Budapest. 833 pp. (in Hungarian)

Sonkoly, J., Tóth, E., Balogh, N., Balogh, L., Bartha, D. ... Török, P. (2022) Introducing PADAPT 1.0, the Pannonian Database of Plant Traits. Journal of Vegetation Science (submitted manuscript)

### Modified Ellenberg – Müller-Dombois life form system

This life form system is an adaptation of Ellenberg & Mueller-Dombois' (1974) system to the vascular plants of Hungary. Categories missing from Hungary (e.g., epiphytes) are not listed. It contains two hierarchical levels: main categories and subcategories. Their meanings are shown in the form of the determination key below:

**Determination key of the main categories**

| | | |
|---|---|---|
| 1 | a) Autotrophic plants ……………………………………………… | 2 |
| | b) Plants not able to photosynthesize …………………………………… | 8 |
| 2 | a) Plants rooting in soil, sediment or floating in water ………..……… | 3 |
| | b) Plants rooting in and taking up water and mineral nutrients from the xylem of other plants …………………………………………………. | **Hemi-parasites (HP)** |
| 3 | a) Self-supporting plants ……………………………………………….. | 4 |
| | b) Plants rooting in the soil, but grow by supporting themselves on others ………………………………………………………………. | **Lianas (L)** |
| | c) Water plants…………………………………………………………... | **Hydrophytes (HH)** |
| 4 | a) Woody plants or herbaceous evergreen perennials………………….... | 5 |
| | b) Herbaceous deciduous plants……………………………………….. | 6 |
| 5 | a) Plants growing taller than 50 cm and shoots do not die back periodically to that height limit……………………………………… | **Phanerophytes (Ph)** |
| | b) Plants whose mature shoot system remains perennially within 50 cm above ground surface or plants that grow taller but whose shoots die back periodically……………………………………………………… | **Chametophytes (Ch)** |
| 6 | a) Annuals, plants whose shoot and root system dies after seed production and which complete their whole life cycle within one year | **Therophytes (Th)** |
| | b) Perennial (including biennial) herbaceous plants with periodic shoot | |

reduction………………………….……………………………………….. 7

7 a) Periodic reduction of the complete above-ground shoot system to storage organs embedded in the soil ………………………………… **Geophytes (G)**

b) Periodic shoot reduction to a remnant shoot system that lies relatively flat on the ground surface …………………………………………… **Hemicryptophytes (H)**

8 a) Growing on living plants…………………………………………... **Parasites (P)**

b) Growing on dead organic matter…………………………………… **Saprophytes (S)**

**Determination key for sub-categories**

**Phanerophytes**

a) Trees, single-stemmed plants with lateral branches………………………………… $\mathbf{P_{scap}}$

b) Shrubs, phanerophytes branched from near the base from the stem……………… $\mathbf{P_{caesp}}$

**Geophytes**

a) Root-budding geophytes…………………………………………………………… $\mathbf{G_{root}}$

b) Bulbous geophytes, arising from bulbs or corms…………………………………... $\mathbf{G_{bulb}}$

c) Rhizomatous geophytes……………………………………………………………. $\mathbf{G_{rhiz}}$

**Therophytes**

a) Spring-green (winter) annuals: germinating from late autumn to spring, flowering in spring or early summer…………………………………………………….…………… $\mathbf{Th_{win}}$

b) Summer-green annuals: germinating from late spring, flowering in summer……… $\mathbf{Th_{sum}}$

**Lianas**

1 a) Woody lianas, including all climbing plants that do not die back periodically to the ground……………………………………………………………………….. **PL**

b) Herbaceous lianas, the above-ground shoot periodically dies back……………… 2

2 a) Annual lianas, they complete their whole life cycle within one year…………… **TL**

b) Perennial herbaceous lianas……………………………………………………… 3

3 a) Periodic reduction of the complete above-ground shoot system to storage organs embedded in the soil…………………………………………………………….. **GL**

b) Periodic shoot reduction to a remnant shoot system that lies relatively flat on the ground surface…………………………………………………………………… **HL**

Within Saprophytes, Parasites, and Hemi-parasites, sub-categories according to life-span and the position of over-wintering buds can be applied similarly to the sub-categories of lianas.

The modified Ellenberg – Müller-Dombois life form system and the categorization of the species was prepared by Zoltán Botta-Dukát.

Data source and citation:

Sonkoly, J., Tóth, E., Balogh, N., Balogh, L., Bartha, D. ... Török, P. (2022) Introducing PADAPT 1.0, the Pannonian Database of Plant Traits. Journal of Vegetation Science (submitted manuscript)

Other references:

Bartha D. (2021) An Annotated and Updated Checklist of the Hungarian Dendroflora. Acta Botanica Hungarica 63(3-4): 227-284.

Horváth, F., Dobolyi, K., Morschhauser, T., Lőkös, L., Karas, L., & Szerdahelyi, T. (1995) Flóra adatbázis 1.2. Taxon-lista és attribútum állomány. Vácrátót: MTA ÖBKI.

Mueller-Dombois, D. & Ellenberg, H. (1974) Aims and methods of vegetation ecology. John Wiley and Sons, New York.
