## Appendix S2 for "PADAPT 1.0 – the Pannonian Database of Plant Traits"

**Supporting Information to the paper Sonkoly, J. et al. Introducing PADAPT 1.0, the Pannonian Database of Plant Traits.** ***Journal of Vegetation Science.***

**Appendix S2. Descriptions and data sources of the attributes in the 'Reproduction' group.**

### Onset and duration of flowering

Onset of flowering: the number denotes the month in which the species most usually starts flowering.

Duration of flowering: duration of the flowering period expressed as number of months.

In case of ferns, the data refer to the time of spore maturation.

Data source and citation:


Király G. (ed.) (2009) Új magyar füvészkönyv. Magyarország hajtásos növényei. Határozókulcsok. [New Hungarian Herbal. The Vascular Plants of Hungary. Identification key.] – Aggteleki Nemzeti Park Igazgatóság, Jósvafő

Sonkoly, J., Tóth, E., Balogh, N., Balogh, L., Bartha, D. ... Török, P. (2022) Introducing PADAPT 1.0, the Pannonian Database of Plant Traits. Journal of Vegetation Science (submitted manuscript)


### Thousand-seed mass (TSM)

Average mass of one thousand seeds of a given species expressed in grams.

In the PADAPT we incorporated published TSM data from Hungary (Schermann 1996; Csontos et al. 2003, 2007; Török et al. 2013, 2016; Balogh et al. 2022).

Data source and citation:

The range of sources that is to be cited is up to the user based on the range of the used data.


Csontos, P., Tamás, J., Balogh, L. (2003) Thousand seed weight records of species from the flora of Hungary, I. Monocotyledonopsida. Studia Botanica Hungarica 34: 121–126.

Csontos, P., Tamás, J., Balogh, L. (2007) Thousand seed weight records of species from the flora of Hungary, II. Dicotyledonopsida. Studia Botanica Hungarica 38: 179–189

Török P., Miglécz T., Valkó O., Tóth K., Kelemen A., Albert Á-J., Matus G., Molnár V. A., Ruprecht E., Papp L., Deák B., Horváth A., Takács A., Hüse B., Tóthmérész B. (2013) New thousand-seed weight records of the Pannonian flora and their application in analysing social behavious types. Acta Botanica Hungarica 55: 429–472.

Török P., Tóth E., Tóth K., Valkó O., Deák B., Kelbert B., Bálint P., Kelemen A., Sonkoly J., Miglécz T., Matus G., Takács A., Molnár V. A., Süveges K., Papp L. , ifj. Papp L., Tóth Z., Baktay B., Málnási Csizmadia G., Oláh I., Peti E., Szalkovszki O., Kiss R., Tóthmérész B.

(2016) New measurements of thousand-seed weight of species in the Pannonian Flora. Acta Botanica Hungarica 58: 187–198.

Schermann, Sz. (1967) Magismeret I–II. – Akadémiai Kiadó, Budapest

**Seed mass category**

Csontos (2001) classified 1676 species of the Hungarian flora into eight seed mass categories based on their thousand-seed mass. The classification was based partly on measured (1479 species) and partly on estimated seed mass data (197 species).

In PADAPT, some of the species not categorised by Csontos (2001) were categorised into the given seed mass categories based on recent publications (Török et al. 2013, Török et al. 2016, Balogh et al. 2022) containing data from seed mass measurements carried out in Hungary. For species that were previously categorised based on estimated values, but for which we now have measured seed mass data, the categorisation based on the measured data was taken into account in PADAPT.

In the case of species that have already been categorised based on measured data but are not categorised in the same seed mass category based on more recent measurements, both classifications are indicated in PADAPT.

In this way, we were able to classify 2214 species (more than 80% of the plant species included in PADAPT) into the appropriate seed mass category.

Ranges of thousand-seed mass values corresponding to the seed mass categories:

| Seed mass category | Thousand-seed mass range |
|---|---|
| 1 | <0.2 g |
| 2 | 0.21 g – 0.50 g |
| 3 | 0.51 g – 1 g |
| 4 | 1.01 g – 2 g |
| 5 | 2.01 g – 4 g |
| 6 | 4.01 g – 10 g |
| 7 | 10.1 g – 50 g |
| 8 | >50 g |

Data source and citation:

Csontos P. (2001) A természetes magbank kutatásának módszerei. Scientia Kiadó, Budapest [in Hungarian]

Sonkoly, J., Tóth, E., Balogh, N., Balogh, L., Bartha, D. ... Török, P. (2022) Introducing PADAPT 1.0, the Pannonian Database of Plant Traits. Journal of Vegetation Science (submitted manuscript)

Other references:

Török P., Miglécz T., Valkó O., Tóth K., Kelemen A., Albert Á-J., Matus G., Molnár V. A., Ruprecht E., Papp L., Deák B., Horváth A., Takács A., Hüse B., Tóthmérész B. (2013) New thousand-seed weight records of the Pannonian flora and their application in analysing social behavious types. Acta Botanica Hungarica 55: 429–472.

Török P., Tóth E., Tóth K., Valkó O., Deák B., Kelbert B., Bálint P., Kelemen A., Sonkoly J., Miglécz T., Matus G., Takács A., Molnár V. A., Süveges K., Papp L. , ifj. Papp L., Tóth Z., Baktay B., Málnási Csizmadia G., Oláh I., Peti E., Szalkovszki O., Kiss R., Tóthmérész B. (2016) New measurements of thousand-seed weight of species in the Pannonian Flora. Acta Botanica Hungarica 58: 187–198.

**Seed bank persistence index**

According to Csontos (2001), seed bank consists of all the naturally occurring seeds that are independent from their mother plants' metabolism and either able to germinate or able to acquire this ability in the future. Several classification systems have been proposed for seed bank types, but Thompson's system (Thompson 1992) is the most well-known and most accepted of them. Thompson described three categories based on the ability of a species' seeds to remain viable in the soil: (i) transient seed bank – seeds remain viable for less than one year, (ii) short-term persistent seed bank – seeds remain viable for more than one year, but less than 5 years, and (iii) long-term persistent seed bank – seeds remain viable for more than 5 years. However, the boundary between the last two categories is often not evident and difficult to detect (Thompson et al. 1993), so in our database we only distinguish transient and persistent seed banks.

To avoid the confounding effect of different climatic and environmental conditions in general, we only considered the results of seed bank studies carried out in Hungary (Csontos 2001, Csontos et al. 2016, Matus et al. 2003, Tóth 2015, Török 2008, and Valkó et al. 2014). Based on the Seed Longevity Index of Bekker et al. (1998), we express the ratio of data indicating a persistent soil seed bank with a value from 0 to 1, where zero means that all available data indicates a transient seed bank, and 1 means that all available data indicates a persistent seed bank.

Data source and citation:

The range of sources that is to be cited is up to the user based on the range of the used data.

Csontos, P. (2001) A természetes magbank kutatásának módszerei. Synbiologia Hungarica, Scientia Kiadó, Budapest.

Csontos, P., Kalapos, T. & Tamás, J. (2016) Comparison of seed longevity for thirty forest, grassland and weed speciesof the Central European Flora: Results of a seed burial experiment. Polish Journal of Ecology 64: 313–326.

Matus, G., Tóthmérész, B. & Papp, M. (2003) Restoration prospects of abandoned species-rich sandy grasslands in Hungary. Applied Vegetation Science 6: 169–178.

Tóth, K. (2015) A magbank szerepe a természetes gyepek diverzitásának fenntartásában és a gyepregenerációban. Doktori disszertáció, Debreceni Egyetem

Török, P. (2008) A magkészlet szerepe mészkerülő gyepek rehabilitációjában. Doktori disszertáció, Debreceni Egyetem

Valkó, O., Török, P., Tóthmérész, B., & Matus, G. (2011) Restoration potential in seed banks of acidic fen and dry-mesophilous meadows: can restoration be based on local seed banks? Restoration Ecology 19: 9–15.

Sonkoly, J., Tóth, E., Balogh, N., Balogh, L., Bartha, D. ... Török, P. (2022) Introducing PADAPT 1.0, the Pannonian Database of Plant Traits. Journal of Vegetation Science (submitted manuscript)

Other references:

Bekker, R. M., Bakker, J. P., Grandin, U., Kalamees, R., Milberg, P., Poschlod, P., ... & Willems, J. H. (1998) Seed size, shape and vertical distribution in the soil: indicators of seed longevity. Functional Ecology 12: 834–842.

Thompson, K. (1992) The functional ecology of seed banks. Seeds: the Ecology of Regeneration in Plant Communities (ed. M. Fenner), pp. 231–258. CAB International, Wallingford, UK.

Thompson, K., Band, S.R. & Hodgson, J.G. (1993) Seed size and shape predict persistence in soil. Functional Ecology 7: 236–241.

**Dispersal strategy**

The concept of dispersal strategies was proposed by Sádlo et al. (2018). Plant species use different dispersal modes (or dispersal syndromes), which are defined by the employed dispersal vector, so anemochory means dispersal by wind, hydrochory is dispersal by water, and zoochory is dispersal by animals, etc. As a single plant species can use not only a single dispersal mode but a set of them, assigning one dispersal mode to a species is most often not feasible. However, certain combinations of dispersal modes occur together in different sets of species, which combinations define dispersal strategies. Nine dispersal strategies have been defined by Sádlo et al. (2018), each of which are named after a genus that represents the strategy well. Dispersal modes involved in each strategy are shown in brackets, with the dominant dispersal modes underlined. Species that are not part of the Czech flora and thus have not been evaluated by Sádlo et al. (2018) were categorised into dispersal strategies following the guideline of Sádlo et al. (2018).

- **Allium type** (<u>autochory</u>, anemochory, endozoochory, epizoochory)
  The most common dispersal strategy, which includes approximately half of the species of the Hungarian flora. Most of the species in this category are dispersal generalists without clear signs of adaptation to anemochory or zoochory.

- **Bidens type** (autochory, epizoochory, endozoochory)

  This strategy is characterised by two main dispersal modes. Despite their clear morphological adaptation for epizoochory, autochory is the most important dispersal mode for the species.

- **Cornus type** (autochory, endozoochory)

  Typically, herbaceous species, shrubs and trees with fleshy fruits belong to this strategy, often from the Rosaceae family. Tree species with heavy and nutrient-rich seeds also belong to this category.

- **Epilobium type** (anemochory, autochory, endozoochory, epizoochory)

  This strategy is characteristic of the species of mesic and dry habitats. The key role of anemochory is evident (several of them are from the Asteraceae family), while the role of autochory is less evident and probably underestimated.

- **Lycopodium type** (anemochory, autochory, endozoochory, epizoochory, hydrochory)

  This strategy relies on very small and light seeds and spores that can easily disperse by not just wind, but also by several other agents. Compared to other dispersal strategies, the role of autochory is quite small for these species.

- **Phragmites type** (anemochory, hydrochory, autochory, endozoochory, epizoochory)

  Species occurring in wetlands and having small diaspores equipped with flying apparatus belong to this strategy. Species in this category most usually do not have vegetative diaspores. Woody species and clonal graminoids and herbs are typical of this strategy.

- **Sparganium type** (autochory, hydrochory, endozoochory, epizoochory)

  This strategy is characteristic of wetland species, but it resembles to the Wolffia type dispersal strategy of aquatic plants. Mostly monocot species having seeds with good buoyancy belong to this category. Vegetative dispersal also plays an important role in this strategy.

- **Wolffia type** (hydrochory, epizoochory, endozoochory)

  Aquatic plants (macrophytes) spreading with fruits, seeds or spores belong to this category, but vegetative dispersal and reproduction, for example by stolons and stem fragmentation, dominates in most of these species.

- **Zea type**

  Species in this category are mostly crops and ornamentals that almost never disperse by generative diaspores and do not have vegetative aboveground diaspores either.

Data source and citation:

Sádlo J., Chytrý M., Pergl J. & Pyšek P. (2018) Plant dispersal strategies: a new classification based on the multiple dispersal modes of individual species. Preslia 90: 1–22.

Sonkoly, J., Tóth, E., Balogh, N., Balogh, L., Bartha, D. ... Török, P. (2022) Introducing PADAPT 1.0, the Pannonian Database of Plant Traits. Journal of Vegetation Science (submitted manuscript)
