## Appendix S3 for "PADAPT 1.0 – the Pannonian Database of Plant Traits"

**Supporting Information to the paper Sonkoly, J. et al. Introducing PADAPT 1.0, the Pannonian Database of Plant Traits.** ***Journal of Vegetation Science.***

**Appendix S3. Descriptions and data sources of the attributes in the 'Kariology' group.**

### Kariology (chromosome number and ploidy level)

An important characteristic of species is the number of chromosomes in their somatic cells. Most of them are diploid, i.e., their somatic cells contain two of each chromosome (1 maternal + 1 paternal). 2n is always used to refer to somatic cells, which means a double chromosome set in the case of diploid cells; 2n=2x, where x is the number of homologous chromosomes.
Ploidy level denotes the number of homologous chromosomes in the cell. Poliploidy, i.e., the multiplication of chromosomes is especially common among plants. Multiplicated chromosome sets in poliploid species result in chromosome numbers 2n=4x, 2n=5x, 2n=6x etc., several types are known. In case of autoploids the full chromosome set is duplicated or multiplicated, generating tetraploids and octoploids, so that chromosome number in somatic cells is 2n=4x and 2n=8x, respectively.

The dataset on chromosome numbers and ploidy levels was prepared by Mária Höhn.

Citation:

Sonkoly, J., Tóth, E., Balogh, N., Balogh, L., Bartha, D. ... Török, P. (2022) Introducing PADAPT 1.0, the Pannonian Database of Plant Traits. Journal of Vegetation Science (submitted manuscript)

Other references:

Bódis J., Biró É., Nagy T., ... Molnár V. A. (2019) Biological flora of Central Europe Himantoglossum adriaticum H. Baumann. Perspectives in Plant Ecology, Evolution and Systematics 40: 125461.

Canne J. M. (1987) Determinations of chromosome numbers in Viola (Violaceae). Canadian Journal of Botany 65: 653–655.

de Castro D. (1949) Novas numeros de cromosomas para o genero Cytisus L. Agronomia Lusitana 11: 85-89.

Ciocarlan V. (2009) Flora ilustrata a Romaniei. Editura CERES.

Dvořak F. (1977) Study of chromosomes of Angiosperms 5. Scripta Fac. Sci. Nat. UJEP Brun., Biol. l: 9–30.

D'Emerico S., Paciola C., Tomasi F. (2000) Contribution to the karyomorphology of some species of the genus Quercus. Silvae Genetica. 49: 243–245.

Dönmez A. A. (2004) The genus Crataegus L. (Rosaceae) with special reference to hybridisation and biodiversity in Turkey. Turkish Journal of Botany 28: 29–37.

Forissier R. (1973) IOPB Chromosome number reports XLVII, A. Löve (ed.). Taxon 24: 143–146. & 671–678.

Forissier R. (1973) Recherches cytotaxonomiques préliminaires sur les genres Lembotropis, Cytisus, Chamaecytisus, Genista et Chamaespartium. Bulletin de la Société Neuchâteloise des Sciences Naturelles 96: 51–65.

Gilot, J. (1965) Contribution à l'étude cytotaxonomique des Genisteae et des Loteae. – Cellule 65: 317–347.

Gregor T., Hand R. (2014) Chromosomenzahlen von Farn- und Samenpflanzen aus Deutschland. Kochia 8: 63–70.

Holubová-Klásková A. (1964) Bemerkungen zur Gliederung der Gattung Cytisus L. s.l. Acta. Universitatis Carolinae Biolgica 2: 1–23.

Kole C. (2007) Forest Trees. Springer.

Kovanda M, (1970) Polyploidy and variation in the Campanula rotundifolia complex. Part II (Taxonomic). 2. Revision of the groups Saxicolae, Lanceolatae and Alpicolae in Czechoslovakia and adjacent regions. Folia Geobotanica & Phytotaxonomica, Praha 5: 171–208.

Majure C. L., Puente R., Griffith P., Judd W. S., Soltis M. P., Soltis E. D. (2012) Phylogeny of Opuntia s.s. (Cactaceae): Clade delineation, geographic origins, and reticulate evolution. American Journal of Botany 99: 847–864.

Löve Á. (1973) IOPB chromosome number reports XLII. Taxon 22: 647-654.

Májovsky J., Murin A. (1987) Karyotaxonomický prehľad flóry Slovenska. Veda, Bratislava

Moore D. M. (1982) Flora Europaea check-list and chromosome index. Cambridge University Press

McKelvey S. D., Sax K. (1933) Taxonomic and cytological relationships of Yucca and Agave. Journal of the Arnold Arboretum 14: 76–81.

Moulanis D., Illies Z. M. (1975) Vergleichende zytologische Untersuchungen der Chromosomenstruktur von Abies borisii-regis Mattf., A. cephalonica Loud, und A. alba Mill. Silvae Genetica 24: 115–118.

Nebel B. R. (1929) Chromosome counts in Vitis and Pyrus. The American Naturalist. 63: 188–189.

Németh Cs. (2015) Sorbus pelsoensis (Sorbus subgenus Tormaria), a new species from the surroundings of Lake Balaton, Hungary. Studia Botanica Hungarica 46: 49–60.

Ohri D., Ahuja M.R. (1990) Giemsa C-banded karyotype in Quercus L. (oak). Silvae Genetica 39: 216–219.

Paule J., Gregor T., Schmidt M., Gerstner E-M., Dersch G., Dressler S., Wesche K., Zizka G. (2017) Chromosome numbers of the flora of Germany – a new online database of georeferenced chromosome counts and flow cytometric ploidy estimates. Plant Systematics and Evolution 303:1123–1129.

Pinkava D. J., McLeod M. G. (1971) Chromosome numbers in some cacti of western North America. Brittonia 23: 171–176.

Rice A., Glick L., Abadi S., Einhorn M., Kopelman N. M., Salman-Minkov A., Mayzel J., Chay O., Mayrose I. (2014) The Chromosome Counts Database (CCDB) – a community resource of plant chromosome numbers. New Phytologist 206: 19–26.

Dos Santos A. C. (1944) Algumas contagens de Cromosomas nos Géneros Genista L. e Cytisus L. Boletim da Sociedade Broteriana 19: 519–522.

Scaltsoyiannes A., Tsaktsira M., Drouzas A. D. (1999) Allozyme differentiation in the Mediterranean firs (Abies, Pinaceae). A first comparative study with phylogenetic implications. Plant Systematics and Evolution 216: 289–307.

Sennikov A. N., Kurtto A. (2017) A phylogenetic checklist of Sorbus s.l. (Rosaceae) in Europe. Memoranda Societatis pro Fauna et Flora Fennica 93: 1–78.

Sochor M., Trávníček B., Király G. (2019) Ploidy level variation in the genus Rubus in the Pannonian Basin and the northern Balkans, and evolutionary implications. Plant Systematics and Evolution 305: 611–626.

Sramkó G., Molnár V. A., Tóth J. P., Laczkó L., Kalinka A., Horváth O., Skuza L., Lukács B.A., Popiela A. (2016) Molecular phylogenetics, seed morphometrics, chromosome number evolution and systematics of European Elatine L. (Elatinaceae) species. PeerJ 4: e2800.

Siljak S. (1971) Basic data on the chromosome complement of the Acer tataricum L. I.er Symposium of Biosystematic in Yougoslavia Proceedings, pp. 139–144.

Zaldos V., Papes D., Brown S. C., Panaus O., Siljak-Yakovlev S. (1998) Genome size and base composition of seven Quercus species: inter and intra-population variation. Genome 41: 162–168.
