## Appendix S4 for "PADAPT 1.0 – the Pannonian Database of Plant Traits"

**Supporting Information to the paper Sonkoly, J. et al. Introducing PADAPT 1.0, the Pannonian Database of Plant Traits. *Journal of Vegetation Science.***

**Appendix S4. Descriptions and data sources of the attributes in the 'Distribution and conservation' group.**

### Phytosociological categorisation

The phytosociological categorisation of species was extracted from the Flora Database (Horváth et al. 1993), but we also categorised some of the previously uncategorised species based on literature data (see below).

#### Soó's phytosociological categorization of the vascular flora of Hungary (COENOS)

Rezső Soó described and systematized the plant communities of Hungary, and established the affiliation, loyalty or indifference of almost all species of the Hungarian flora to coenotaxa. In the Flóra database, the basis for the classification is the phytosociological system of Soó (1968, 1980), supplemented by data from Simon (1992).

Species with a single phytosociological emphasis are associated with a single coenotaxon. Of the species with double or triple emphasis, species with a definite main emphasis were assigned to a category according to their dominant coenotaxon, species with a double emphasis were assigned to double categories, while species "loyal" to more than two coenotaxa were assigned to indifferent (I) or more or less indifferent (H) categories.

Categorisation requires highlighting salient features but losing more nuanced information, so the system can only consider species emphasis in fidelity, which can vary considerably between regions. The phytosociological system is hierarchical, which can give a sense not only of the coenologic emphasis but also of the breadth of species' coenologic behaviour.

In PADAPT, some of the previously uncategorised species have also been categorised based on recent literature data.

Categories and coding of Soó's system:

1 – LEMNO-POTAMEA

11 – Lemnetea

111 – Hydrocharietalia

1111 – Lemnion minoris

*111101 – Lemno-Spirodeletum*

*111102 – Wolffietum arrhizae*

*111103– Salvinio-Spirodeletum*

1112 – Hydrocharition

*111201 – Lemno-Utricularietum*

*111202 – Hydrochari-Stratiotetum*

*111203 – Spirodelo-Aldrovandetum*

12 – Potamogetonetea

121 – Potametalia

1211 – Ranunculion fluitantis

*121101 – Ranunculetum fluitantis*

*121102 – Ranunculetum trichophylli-callitrichetum*

*121103 – Hottonietum palustris*

1212 – Potamogetonion

*121201 – Elodeetum*

*121202 – Myriophyllo-Potamogetonetum*

*121203 – Potamogetonetum lucentis*

*121204 – Najadetum minoris*

1213 – Nymphaeion

*121301 – Potamogetonetum natantis*

*121302 – Nymphaeetum albo-luteae*

*121303 – Nymphoidetum peltatae*

*121304 – Trapetum natantis*

122 – Ruppietalia

1221 – Ruppion maritimae

*122101 – Parvipotameto-Zannichellietum*

123 – Utricularietalia

1231 – Sphagno-Utricularion

2 – CYPERO-PHRAGMITEA

21 – Phragmitetea & Molinio-Juncetea (resp. „Populetalia")

22 – Phragmitetea

221 – Phragmitetalia

2211 – Phragmition australis

*221101 – Scirpo-Phragmitetum*

*221102 – Sparganietum erecti*

*221103 – Glycerietum maximae*

*221104 – Cladietum marisai*

*221105 – Rorippo-Oenanthetum*

222 – Bolboschoenetalia

2221 – Bolboschoenion maritimi continentale

*222101 – Bolboschoenetum maritimi*

*222102 – Schoenoplectetum litoralis*

*222103 – Bolboschoeno-Phragmitetum*

223 – Nasturtio-Glycerietalia

2231 – Glycerio-Sparganion

*223101 – Sparganio-Glycerietum fluitantis*

224 – Magnocaricetalia

2241 – Magnocaricion & Molinietalia

2242 – Magnocaricion elatae

22421 – Caricion rostratae

*224211 – Caricetum elatae*

*224212 – Caricetum appropinquatae*

*224213 – Carici-Menyanthetum*

*224214 – Schoenoplecto-Juncetum maritimi*

22422 – Caricion gracilis

*224221 – Carici-Typhoidetum*

*224222 – Caricetum acutiformis-ripariae*

*224223 – Caricetum vulpinae*

23 – Isoëto-Nanojuncetea

231 – Nanocyperetalia

2311 – Elatini-Eleocharition ovatae & Potamogetonion (resp. Oryzion)

2312 – Elatini-Eleocharition ovatae & Aphanion (resp. Thero-Airion)

2313 – Elatini-Eleocharition ovatae

*231301 – Eleochari-Caricetum bohemicae*

*231302 – Cypero-Juncetum bufonii*

*231303 – Dichostyli-Gnaphalietum uliginosi*

*231304 – Eleochari aciculari-Schoenoplectetum supini*

2314 – Verbenion supinae

24 – Montio-Cardaminetea

241 – Montio-Cardaminetalia

2411 – Cardamino-Montion

*241101 – Cardaminetum amarae*

2412 – Cratoneurion commutati

*241201 – Caricetum lepidocarpae-Cratoneuretum*

3 – OXYCOCCO-CARICEA

31 – Scheuchzerio-Caricetea nigrae & Magnocaricion

32 – Scheuchzerio-Caricetea nigrae

321 – Scheuchzerio-Caricetalia nigrae

3211 – Caricion lasiocarpae

*321101 – Carici lasiocarpae-Sphagnetum*

3212 – Caricion canescenti-nigrae

*321201 – Carici echinatae-Sphagnetum*

3213 – Rhynchosporion albae

33 – Oxycocco-Sphagnetea

331 – Sphagnetalia magellanici

3311 – Sphagnion magellanici

*331101 – Eriophoro vaginati-Sphagnetum recurvi*

4 – MOLINIO-ARRHENATHEREA & CHENOPODIO-SCLERANTHEA

5 – MOLINIO-ARRHENATHEREA

51 – Molinio-Juncetea & Scheuchzerio-Caricetea nigrae

52 – Molinio-Juncetea & Puccinellietalia

53 – Molinio-Juncetea

531 – Caricetalia davallianae' & Molinion

532 –Caricetalia davallianae

5321 – Caricion davallianae

*532101 – Valeriano dioicae-Caricetum davallianae*

*532102 – Schoenetum nigricantis*

*532103 – Juncetum subnodulosi*

*532104 – Seslerietum uliginosae*

5322 – Eriophorion gracilis

*532201 – Carici flavae-Eriophoretum*

533 – Molinietalia (resp. Molinio-Juncetea) & Magnocaricion

534 – Molinietalia & Festucetalia valesiacae (Brometalia in the west)
535 – Molinietalia & Festucion vaginatae
536 – Molinietalia
5361 – Molinion
*536101 – Junco-Molinietum*
*536102 – Succiso-Molinietum*
*536103 – Molinio-Salicetum rosmarinifoliae*
5362 – Deschampsion caespitosae & Puccinellietalia (spec. Beckm.ion & Juncion gerardi)
5363 – Deschampsion caespitosae
*536301 – Deschampsietum caespitosae*
*536302 – Agrostetum albae*
5364 – Alopecurion pratensis
*536401 – Alopecuretum pratensis*
*536402 – Festucetum pratensis*
5365 – Filipendulo-Petasition
*536501 – Petasitetum hybridi*
*536502 – Filipendulo ulmariae-Geranietum palustris*
*536503 – Angelico-Cirsietum oleracei*
54 – Arrhenatheretea
541 – Arrhenatheretalia & Alopecurion pratensis resp. Deschampsion caespitosae
542 – Arrhenatheretalia
5421 –Arrhenatherion elatioris
*542101 – Pastinaco-Arrhenatheretum*
*542102 – Alopecuro-Arrhenatheretum*
*542103 – Anthyllido-Festucetum rubrae*
5422 – Triseto-Polygonion bistortae
*542201 – Trisetetum flavescentis*
5423 – Cynosurion cristati
*542301 – Lolio-Cynosuretum*
55 – Nardo-Callunetea & Molinio-Juncetea
56 – Nardo-Callunetea & Arrhenatheretea
57 – Nardo-Callunetea (often also Pino-Quercetalia)
571 – Nardetalia

5711 – Nardo-Agrostion tenuis

*571101 – Agrostietum tenuis*

*572102 – Festuco ovinae-Nardetum*

572 – Vaccinio-Genistetalia

5721 – Calluno-Genistion

*572101 – Luzulo albidae-Callunetum*

6 – PUCCINELLIO-SALICORNEA

61 - Thero-Salicornietea

611 – Thero-Salicornietalia

6111 –Thero-Salicornion

*611101 – Salicornietum prostratae*

6112 – Thero-Suaedion

*611201 – Suaedetum pannonicae*

612 – Crypsidetalia aculeatae

6121 – Cypero-Spergularion

*612101 – Crypsydetum aculeatae*

*612102 – Acorelletum pannonicae*

62 – Festuco-Puccinellietea

621 – Puccinellietalia

6211 – Puccinellion limosae

*621101 – Puccinellietum limosae*

*621102 – Pholiuro-Plantaginetum*

*621103 – Camphorosmetum annuae*

6212 – Puccinellion peisonis

*621201 – Puccinellietum peisonis*

*621202 – Lepidio-Puccinellietum limosae*

*621203 – Lepidio-Camphorosmetum annuae*

6213 – Juncion gerardii

*621301 – Agrostio-Caricetum distantis*

6214 – Beckmannion eruciformis

*621401 – Agrostio-Alopecuretum pratensis*

*621402 – Agrostio-Glycerietum poiformis*

*621403 – Agrostio-Beckmannietum*

622 – Artemisio-Festucetalia pseudovinae & Festucetalia valesiacae

623 – Artemisio-Festucetalia pseudovinae

6231 – Festucion pseudovinae

*623101 – Artemisio-Festucetum pseudovinae*

*623102 – Peucedano-Asteretum punctati*

7 – SEDO-CORYNEPHOREA

71 – Koelerio-Corynephoretea

711 – Corynephoretalia

7111 – Corynephorion

*711101 – Corynephoretum canescentis*

7112 – Thero-Airion

*711201 – Filagini-Vulpietum pannonicum*

*711202 – Thymo-Festucetum pseudovinae*

*711203 – Festuco ovinae-Rhacomitrietum*

*711204 – Festuco ovinae-Polytrichetum*

72 – Sedo-Polypodietea

721 – Alysso-Sedetalia

7211 – Alysso-Sedion

*721101 – Grimmio-Sedetum albi-sexangularis*

*721102 – Geranio rotundifolio-Sedetum albi*

722 – Hypno-Polypodietalia

7221 – Polypodion

*722101 – Ctenidio-Polypodietum*

*722102 – Hypno-Polypodietum*

8 – FESTUCO-BROMEA & Koelerio-Corynephoretea

9 – FESTUCO-BROMEA

91 – Festucetea vaginatae

911 – Festucetalia vaginatae & Festucetalia valesiacae

912 – Festucetalia vaginatae

9121 – Bromion tectorum

*912101 – Brometum tectorum*

*912102 – Brometum secaletosum*

9122 – Festucion vaginatae & Festucion rupicolae

9123 – Festucion vaginatae

*912301 – Festucetum vaginatae*

*912302 – Festuco- Corynephoretum*

92 – Festuco-Brometea & Arrhenatheretea

94 – Festuco-Brometea (or Festucetalia valesiacae previously, nowadays mostly Chenopodietea)

95 – Festuco-Brometea (resp. Festucetalia val.) & Quercetea pubescenti-petraeae

96 – Festuco-Brometea

961 – Brometalia erecti

9611 – Cirsio-Brachypodion

*961101 – Lino tenuifolio-Brachypodietum pinnati*

*961102 – Galio boreali-Brachypodietum pinnati*

962 – Festucetalia valesiacae

9621 – Seslerio-Festucion pallentis & Asplenio-Festucetum pallentis

9622 – Asplenio-Festucion pallentis

*962201 – Minuartio-Festucetum pseudodalmaticae et poétosum scabrae*

*962202 – Inulo-Festucetum pseudodalmaticae*

9623 – Bromo-Festucion pallentis

*962301 – Seseli leucospermo-Festucetum pallentis*

*962302 – Stipo-Festucetum pallentis*

*962303 – Seslerietum sadlerianae*

*962304 – Festuco pallenti-Brometum erecti-pannonici*

*962305 – Sedo sopianae-Festucetum dalmaticae*

9624 – Seslerio-Festucion pallentis

*962401 – Campanulo-Festucetum pallentis*

9625 – Festucion rupicolae

*962501 – Chrysopogono-Caricetum humilis*

*962502 – Cleistogeno-Festucetum rupicolae*

*962503 – Pulsatillo-Festucetum rupicolae*

*962504 – Medicagini-Festucetum valesiacae*

*962505 – Salvio-Festucetum rupicolae*

*962506 – Astragalo-Festucetum rupicolae*

9626 – Cynodonto-Festucion rupicolae-pseudovinae

*962601 – Potentillo arenariae-Festucetum pseudovinae*

*962602 – Cynodonto-Poétum angustifoliae*

*962603 – Achilleo-Festucetum pseudovinae*

*962604 – Cynodonto-Festucetum pseudovinae*

9627 – Danthonio-Stipion tirsae

*962701 – Campanulo-Stipetum tirsae*

9628 – Artemisio-Kochion

*962801 – Agropyro pectinati-Kochietum prostratae*

A – CHENOPODIO-SCLERANTHEA

A1 – Secalietea

A11 – Aperetalia

A111 – Aphanion

*A111101 – Setario-Digitarietum*

A12 – Lolio-Linetalia

A121 – Lolio remoto-Linion

*A12101 – Lolio temulenti-Linetum*

A13 – Secalietalia

A131 – Caucalydion platycarpos

*A13101 – Setario-Stachietum*

A132 – Trifolio-Medicaginion

A14 – Eragrostetalia

A141 – Consolido- & Tribulo-Eragrostion & Festucion vaginatae

A142 – Consolido- & Tribulo-Eragrostion

A143 – Consolido-Eragrostion minoris

*A14301 – Consolido orientali-Stachyetum annuae*

*A14302 – Amarantho-Chenopodietum albi*

*A14303 – Convolvulo-Portulacetum*

A144 – Tribulo-Eragrostion minoris

*A14401 – Digitario-Portulacetum*

A15 – Oryzetalia

A151 – Oryzion sativae

*A15101 – Echinochloo-Oryzetum sativae*

A2 – Chenopodietea & Festucetalia valesiacae

A3 – Chenopodietea (incl. Sisymbrietalia & Sisymbrion)

A31 – Sisymbrietalia

A311 – Sisymbrion

*A31101 – Rorippo austriacae-Hordeetum murini*

*A31102 – Atriplicetum tataricae*

*A31103 – Malvetum neglectae*

A312 – Convolvulo-Agropyrion repentis

*A31201 – Agropyro-Convolvuletum arvensis*

A32 – Onopordietalia (incl. Onopordion)

A321 – Onopordion acanthii

*A32101 – Onopordetum acanthii*

*A32102 – Xanthietum spinosi*

A4 – Artemisietea

A41 – Artemisietalia

A411 – Arction lappae

*A41101 – Conietum maculati*

*A41102 – Arctio-Ballotetum nigrae*

A5 – Galio-Urticetea

A51 – Calystegietalia

A511 – Alliarion petiolatae

*A51101 – Chaerophylletum bulbosi*

*A51102 – Rudbeckio-Solidaginetum*

A512 – Calystegion sepium (resp. „Populetalia")

*A51201 – Glycyrrhizetum echinatae*

A6 – Bidentetea & Elatini-Eleocharition ovatae (resp. Agrostion)

A7 – Bidentetea tripartitae

A71 – Bidentetalia (incl. Bidention)

A711 – Bidention tripartitae

*A71101 – Bidentetum*

A712 – Chenopodion rubri

*A71201 – Dichostyli-Chenopodietum rubri*

*A71202 – Echinochloo-Polygonetum lapathifolii*

A8 – Plantaginetea

A81 – Plantaginetalia (incl. Agropyro-Rumicion crispi)

A811 –Polygonion avicularis

*A81101 –Lolio-Plantaginetum*

*A81102 – Sclerochloo-Polygonetum avicularis*

*A81103 – Juncetum tenuis*

A812 – Agropyro-Rumicion crispi

*A81201 – Lolio-Potentilletum anserinae*

*A81202 – Lolio-Alopecuretum*

A9 – Epilobietea angustifolii

A91 – Epilobietalia angustifolii

A911 – Epilobion angustifolii & Atropion bella-donnae

A912 – Epilobion angustifolii

*A91201 – Senecioni-Epilobietum angustifolii*

*A91202 – Calamagrostietum epigeii*

A913 – Atropion bella-donnae

*A91301 – Atropetum bella-donnae*

*A91302 – Molinietum litoralis*

A92 – Sambucetalia

A921 – Sambuco-Salicion capreae

*A92101 – Fragario-Rubetum*

*A92102 – Sambucetum racemosae*

B – QUERCO-FAGEA & MOLINIO-ARRHENATHEREA

C – QUERCO-FAGEA (spec. Fagetalia) & Epilobietea

D – QUERCO-FAGEA with an emphasis on Quercetea

E – QUERCO-FAGEA

E1 – Salicetea purpureae

E11 – Salicetalia purpureae

E111 – Salicion eleagni

*E11101 – Myricario-Epilobetum*

*E11102 – Hippophaë-Salicetum elaeagni*

E112 – Salicion triandrae

*E11201 – Salicetum purpureae*

*E11202 – Salicetum triandrae*

E113 – Salicion albae (resp."Populetalia")

*E11301 – Salicetum albae-fragilis*

E2 – Alnetea glutinosae & Alno-Padion

E3 – Alnetea glutinosae

E31 – Alnetalia glutinosae

E311 – Alnion glutinosae

*E31101 – Thelypteridi-Alnetum*

*E31102 – Dryopteridi-Alnetum*

*E31103 – Fraxino pannonicae-Alnetum*

E312 – Salicion cinereae

*E31201 – Calamagrostio-Salicetum cinereae*

*E31202 – Salici cinereae-Sphagnetum recurvi*

*E31203 – Salici pentandrae-Betuletum pubescentis*

E4 – Querco-Fagetea

E41 – Fagetalia

E411 – Alno-Padion

*E41101 – Fraxino pannonicae-Ulmetum*

E412 – Alnion glutinosae-incanae

*E41201 – Carici remotae-Fraxinetum*

*E41202 – Carici brizoidis-Alnetum*

*E41203 – Carici acutiformis-Alnetum*

*E41204 – Aegopodio-Alnetum*

E413 – Fagion medio-europaeum

*E4131 – Asperulo-Fagion*

*E41311 – Aconito-Fagetum*

*E41312 – Melittio-Fagetum*

*E41313 – Abieti-Fagetum*

E4132 – Cephalanthero-Fágion (& Orno-Fagetum)

*E41324 – Seslerio hungaricae-Fagetum*

*E41325 – Tilio-Sorbetum*

E4133 – Tilio-Acerion & Cephalanthero-Fagion

E4134 – Tilio-Acerion

*E41346 – Phyllitidi & Parietario Aceretum*

E4135 – Carpinion betuli

*E41357 – Querco petraeae-Carpinetum*

*E41358 – Querco robori-Carpinetum*

*E41359 – Aceri campestri-Quercetum petraeae-roboris*

E414 – Fagion illyricum

*E41401 – Fraxino pannonicae-Carpinetum*

*E41402 – Helleboro dumetoro-Carpinetum*

*E41403 – Asperulo taurinae-Carpinetum*

*E41404 – Vicio oroboidi-Fagetum*

*E41405 – Helleboro odoro-Fagetum*

*E41406 – Tilio tomentosae-Fraxinetum*

E415 – Fagion dacicum

E42 – Pino -Quercetalia

E421 – Castaneo-Quercion

*E42101 – Castaneo-Quercetum*

E422 – Genisto germanicae-Quercion

*E42201 – Luzulo-Quercetum subcarpaticum*

*E42202 – Genisto pilosae-Quercetum petraeae*

*E42203 – Sorbo-Quercetum petraeae*

E423 – Pino-Quercion

*E42301 – Pino-Quercetum*

E424 – Deschampsio-Fagion

*E42401 – Luzulo-Fagetum*

E5 – Quercetea pubescenti-petraeae & Pino-Quercetalia

E6 – Quercetea pubescenti-petraeae

E61 – Orno-Cotinetalia

E611 – Orno-Cotinion

*E61101 – Cotino-Quercetum pubescentis*

*E61102 – Fago-Ornetum*

*E61103 – Orno-Quercetum pubescenti-cerris*

E612 – Quercion farnetto

*E61201 – Potentillo micranthae-Quercetum*

E62 – Quercetalia pubescentis

E621 – Quercion petraeae & Aceri tatarico-Quercion

E622 – Quercion petraeae

*E62201 – Quercetum petraeae-cerris*

E623 – Aceri tatarico-Quercion & Festucion rupicolae

E624 – Aceri tatarico-Quercion

*E62401 – Ceraso-Quercetum pubescentis*

*E62402 – Corno-Quercetum pubescentis-petraeae*

*E62403 – Waldsteinio-Spiraeetum mediae*

*E62404 – Tilio-Fraxinetum*

*E62405 – Mercuriali-Tilietum*

*E62406 – Aceri tatarico-Quercetum pubescenti-roboris*

*E62407 – Dictamno-Tilietum cordatae*

*E62408 – Festuco rupicolae-Quercetum roboris*

*E62409 – Festuco-Populo-Quercetum roboris*

*E6240A – Junipero-Populetum albae*

*E6240B – Galatello-Quercetum roboris*

*E6240C – Convallario-Quercetum roboris*

E63 Prunetalia

E631 Prunion spinosae & Cerasion fruticosae

E632 Cerasion fruticosae

*E63201 Amygdaletum nanae*

*E63202 Crataego-Cerasetum fruticosae*

E633 Prunion spinosae

*E63301 Primo spinosae-Crataegetum*

E634 Corylion

*E63401 Coryletum avellanae*

*E63402 Solidagini-Cornetum*

F – ABIETI-PICEEA

F1 Erico-Pinetea

F11 Erico-Pinetalia

F111 Erico-Pinion

*F11101 Chamaebuxo-Pinetum*

*F11102 Lino flavae-Pinetum*

F12 Pulsatillo-Pinetalia

F121 – Festuco vaginatae-Pinion

*F12101 – Festuco vaginatae-Pinetum*

F2 Vaccinio-Piceetea

F21 Vaccinio-Piceetalia

F211 Abieti-Piceion

*E21101 Bazzanio-Abietetum praealpinum*

G – SILVAE CULTAE

H – More or less indifferent species

I – Indifferent species

**Borhidi's phytosociological categorization of the vascular flora of Hungary (COENOB)**

Attila Borhidi (1995) applied Ellenberg's work to the Hungarian conditions to create a system of Hungarian plant associations.

Overview of the Hungarian plant associations within the framework of the Central European plant associations:

1. Water, swamp and moor vegetation

1.1 Lemnetea

1.1.1 Lemnetalia

1.1.1.1 Lemnion

Lemnetum gibbae

Lemnetum minoris

Lemno-Spirodeletum

Salvinio-Spirodeletum

1.1.2 Hydrocharetalia

1.1.2.1 Hydrocharition

Hydrochari-Stratiotetum

Lemno-Utricularietum

1.2 Utricularietea

1.2.1 Utricularietalia

1.2.1.1 Sphagno-Utricularion

Aldrovando-Utricularietum minoris

Riccietum fluitantis

Spirodelo-Aldrovandetum

1.3 Potamogetonetea

1.3.1 Potamogetonetalia

1.3.1.1 Potamogetonion

Elodeetum

Myriophyllo-Potametum

Najadetum minoris

Parvipotameto-Zannichellietum

Potamogetonetum lucentis

Potamogetonetum pectinati

1.3.1.2 Nymphaeion

Hippuridetum

Nymphaeetum albo-luteae

Polygonetum natantis

Nymphoidetum peltatae

Trapetum natantis

1.3.1.3 Batrachion fluitantis

Callitrichetum cophocarpae

Elatinetum triandrae

Hottonietum palustris

Ranunculetum aquatilis

Ranunculetum fluitantis

Ranunculetum polyphylli

1.4 Litorelletea

1.4.1 Litorelletalia

1.4.1.1 Litorellion

Ranunculo flammulae-Gratioletum

1.5 Phragmitetea

1.5.1 Phragmitetalia

1.5.1.1 Phragmition

Acoretum

Cladietum marisci

Glycerietum maximae

Rorippo-Oenanthetum

Scirpo-Phragmitetum

Schoenoplectetum lacustris

Typhetum angustifoliae

Typhetum latifoliae

1.5.1.2 Eleochari-Sagittarion

Alismato-Eleocharitetum

Butometum umbellatae

1.5.1.3 Glycerio-Sparganion

Glycerietum plicatae

Leersietum

Rorippo-Typhoidetum

Sparganio-Glycerietum fluitantis

1.5.1.4 Magnocaricion

1.5.1.4.1 Caricion rostratae

Calamagrostetum canescentis

Caricetum appropinquatae

Caricetum elatae

Caricetum paniculatae

Carici-Calamagrostetum neglectae

Carici-Menyanthetum

Schoenoplecto-Juncetum maritimi

Caricion gracilis

Caricetum acutiformis-ripariae

Caricetum gracilis

Caricetum vesicariae
Caricetum vulpinae
Carici-Typhoidetum

1.6 Montio-Cardaminetea
- 1.6.1 Montia-Cardaminetalia
  - 1.6.1.1 Montio-Cardaminion
    - Bryetum schleicheri
    - Chrysosplenio-Cardaminetum amarae
  - 1.6.1.2 Cratoneurion commutati
    - Carici lepidocarpae-Cratoneuretum

1.7 Scheuchzerio-Caricetea nigrae
- 1.7.1 Scheuchzerio-Caricetalia
  - 1.7.1.1 Rhynchosporion
    - Rhynchosporetum albae
  - 1.7.1.2 Caricion lasiocarpae
    - Caricetum appropinquatae-stellatae
    - Caricetum canescentis-nigrae
    - Carici echinatae-Sphagnetum
    - Carici lasiocarpae-Sphagnetum
- 1.7.2 Tofieldietalia
  - 1.7.2.1 Caricion davallianae
    - Cladio-Schoenetum
    - Orchido-Schoenetum
    - Schoeno-Seslerietum
    - Valeriano-Caricetum davallianae
- 1.7.3 Caricetalia nigrae
  - 1.7.3.1 Caricion nigrae (fuscae)
    - Junceto-Caricetum nigrae

1.8 Oxycocco-Sphagnetea
- 1.8.1 Sphagnetalia magellanici
  - 1.8.1.1 Sphagnion magellanici (fusci)
    - Eriophoro vaginati-Sphagnetum

2. Coastal and salt swamp vegetation

- 2.1 Zosteretea
- 2.2 Ruppietea
  - 2.1.1 Ruppietalia
    - 2.1.1.1 Ruppion maritimae
      - Parvipotameto-Zannichellietum
      - Ranunculetum aquatilis-polyphylli
- 2.3 Spartinetea
- 2.4 Thero-Salicornietea
  - 2.4.1 Thero-Salicornietalia
    - 2.4.1.1 Salicornion dolichostachyae
    - 2.4.1.2 Salicornion ramosissimae
      - Salicornietum prostratae
      - Salsoletum sodae
      - Suaedetum pannonicae
  - 2.4.2 Crypsidetalia
    - 2.4.2.1 Cypero-Spergularion
      - Acorelletum pannonici
      - Chenopodietum urbici
      - Crypsidetum aculeatae
      - Heleocloetum alopecuroidis
      - Heleochloetum schoenoidis
- 2.5 Saginetea
- 2.6 Asteretea tripolii
- 2.7 Honckenyo-Eymetea
- 2.8 Cakiletea
- 2.9 Ammophiletea
- 2.10 Bolboschoenetea
  - 2.10.1 Bolboschoenetalia
    - 2.10.1.1 Bolboschoenion maritimi
      - Bolboschoenetum maritimi
      - Bolboschoeno-Phragmitetum
      - Polygono-Bolboschoenetum

Schoenoplectetum litoralis

Schoenoplectetum tabemaemontani

3. Disturbed and secondary vegetation

3.1 Isoeto-Nanojuncetea

3.1.1 Cyperetalia fusci

3.1.1.1 Nanocyperion

Cyperetum fusci

Cypero-Juncetum bufonii

Elatini-Lindernietum procumbentis

Eleochari-Schoenoplectetum supini

Ranunculo lateriflori-Limoselletum

3.1.1.2 Elatini-Eleocharion ovatae

Dichostyli-Gnaphalietum uliginosae

Eleochari-Caricetum bohemicae

Lindemio-Eleocharetum ovatae

3.1.1.3 Heleochloo-Cyperion

Dichostyli-Heleochloetum alopecuroidis

Lythretum hyssopifolii-tribracteati

Lythro-Gnaphalietum luteo-albi

3.1.1.4 Verbenion supinae

Astragalo-Chlorocyperetum glomerati

Centunculo-Radioletum linoidis

Heliotropio-Verbenetum supinae

Lythro-Pulicarietum

3.2 Bidentetea

3.2.1. Bidentetalia

3.2.1.1 Bidention tripartitae

Eleochloo-Bidentetum tripartitae

Polygono lapathifolio-Bidentetum

Stachydi-Bidentetum tripartitae

Xanthio strumarii-Bidentetum

3.2.1.2 Chenopodion rubri

Chenopodietum glauci-rubri

Dichostylidi-Chenopodietum rubri

Echinochloo-Polygonetum lapathifolii

Echinochloo-Setarietum pumilae

3.3 Chenopodietea

3.3.1 Polygono-Chenopodietalia

3.3.1.1 Fumario-Euphorbion

3.3.1.2 Spergulo-Oxalidion

3.3.2 Eragrostietalia

3.3.2.1 Eragrostion

Amarantho-Chenopodietum albi

3.3.2.2 Digitario-Setarion

3.3.2.3 Consolido-Eragrostion minoris

Consolido orientali-Stachyetum annuae

Convolvulo-Portulacetum

3.3.2.4 Tribulo-Eragrostion minoris

Digitario-Portulacetum

Hibisco-Eragrostietum minoris

Tribulo-Tragetum

Vicio-Eragrostietum minoris

Vicio-Polygonetum arenarii

3.3.2.5 Matricario-Chenopoclion albi

Matricario-Atriplicetum litoralis

3.3.3 Sisymbrietalia

3.3.3.1 Sisymbrion officinalis

Bromo arvensi-Hordeetum murini

Chenopoclio vulvariae-Urticetum urentis

Descurainietum sophiae

Lappulo-Cynoglossetum

Malvetum neglectae

Malvetum pusillae

Rorippo austriacae-Hordeetum murini

3.3.3.2 Salsolion ruthenicae

Atriplicetum tatarici

3.4 Secalietea

3.4.1 Secalietalia

3.4.1.1 Caucalion lappulae

Setario-Stachyetum annuae

3.4.1.2 Trifolio-Medicaginion

Plantagini-Medicaginetum

3.4.2 Aperetalia

3.4.2.1 Aphanion

Aphani-Matricarietum chamomillae

Rorippo-Setarietum

Sclerantho-Trifolietum arvensis

Setario-Digitarietum

Spergulo-Aperetum

3.4.3 Lolio-Linetalia

3.4.3.1 Lolio-Linion

Lolio temulenti-Linetum

3.5 Artemisietea

3.5.1 Artemisietalia

3.5.1.1 Arction lappae

Arctio-Ballotetum nigrae

Conietum maculati

Lycietum barbarum

Pruno-Ballotetum nigrae

Sambucetum ebuli

Tanaceto-Artemisietum

3.5.2 Calystegietalia

3.5.2.1 Calystegion sepium

Asteri-Rubetum caesii

Bidenti-Calystegietum

Cuscuto-Calystegietum

Glycyrrhizetum echinatae

lmpatienti-Calystegietum

Rudbeckio-Solidaginetum

3.5.3 Glechometalia

3.5.3.1 Aegopodion podagrariae

Chaerophylletum aromatici

Chaerophylletum bulbosi

Urtico-Aegopodietum

3.5.3.2 Alliarion

Alliario-Chaerophylletum temuli

Arctietum nemorosi

Cephalarietum pilosae

Chelidonio-Alliarietum

Epilobio montani-Geranietum robertiani

Eupatorietum cannabini

Torilidetum japonicae

3.5.4 Onopordetalia

3.5.4.1 Onopordion acanthii

Onopordetum acanthii

Xanthietum spinosi

3.5.4.2 Dauco-Melilotion

Berteroetum incanae

Echio-Melilotetum albi

Lactucetum salignae

3.6 Agropyretea

3.6.1 Agropyretalia repentis

3.6.1.1 Convolvulo-Agropyrion

Agropyro-Convolvuletum arvensis

Aristolochio-Convolvuletum arvensis

Bromo japonici-Aristolochietum

Cardarietum drabae

Junco-Tussilaginetum

Stellario mediae-Mercurialietum annuae

3.6.1.2 Artemisio-Agropyrion

Artemisio absinthii-Agropyretum intennedii

Festuco-Agropyretum intermedii

3.7 Plantaginetea

3.7.1 Plantaginetalia majoris

3.7.1.1 Polygonion avicularis

Amarantho deflexo-Polygonetum

Eragrosti-Polygonetum

Euphorbio supinae-Polygonetum avicularis

Juncetum tenuis

Lolio-Plantaginetum Poetum annuae

Sagino-Bryetum argenteae

Sclerochloo-Polygonetum avicularis

3.8 Agrostietea stoloniferae

3.8.1 Agrostietalia stoloniferae

3.8.1.1 Agropyro-Rumicion crispi

Amarantho deflexo-Polygonetum

Blysmo-Juncetum compressi

Dactylido-Festucetum arundinaceae

Juncetum effusi

Junco-Menthetum longifoliae

Lolio-Alopecuretum

Lolio-Potentilletum anserinae

Ranunculetum repentis

Rorippo austriacae-Agropyretum repentis

Rorippo sylvestri-Agrostietum stoloniferae

Rumici-Alopecuretum geniculati

Trifolio fragifero-Cynodontetum

Trifolio repenti-Lolietum

3.9 Oryzetea

3.9.1 Oryzetalia

3.9.1.1 Oryzion sativae

Echinochloo-Oryzetum sativae

4. Montane rock vegetation and alpine grasslands

5. Antropo-zoogenous heath, grasslands and pastures

- 5.1 Nardo-Callunetea
  - 5.1.1 Nardetalia
    - 5.1.1.1 Nardion
    - 5.1.1.2 Violion caninae
    - 5.1.1.3 Juncion squarrosi
    - 5.1.1.4 Festucion variae
    - 5.1.1.5 Nardo-Agrostion tenuis
      - Agrostietum strictae-tenuis
      - Festuco ovinae-Nardetum
      - Festuco tenuifoliae-Agrostietum tenuis
  - 5.1.2 Calluno-Ulicetalia
    - 5.1.2.1 Calluno-Genistion
      - Luzulo luzuloidis-Callunetum
- 5.2 Sedo-Scleranthetea
  - 5.2.1 Sedo-Scleranthetalia
    - 5.2.1.1 Sedo-Scleranthion
    - 5.2.1.2 Alysso-Sedion
      - Geranio rotundifolio-Sedetum albi
      - Grimmio-Sedetum albi-sexangularis
      - Hypno-Sedetum
      - Sedo sexangulari-Allietum montani
    - 5.3.1.3 Seslerio-Festucion pallentis
      - Seslerio-Festucetum pallentis
  - 5.2.2 Corynephoretalia
    - 5.2.2.1 Corynephorion
      - Thymo angustifolio-Corynephoretum
  - 5.2.3 Festuco-Sedetalia
    - 5.2.3.1 Koelerion glaucae
    - 5.2.3.2 Sileno conicae-Cerastion semidecandri
  - 5.2.4 Thero-Airetalia
    - 5.2.4.1 Thero-Airion

- Festuco ovinae-Polytrichetum
- Festuco ovinae-Rhacomitrietum
- Festuco-Thymetum serpylli
- Filagini-Vulpietum
- Melampyro-Rhacomitrietum

5.2.5 Hypno-Polypodietalia

5.2.5.1 Polypodion

- Hypno-Polypodietum

5.3 Festuco-Brometea

5.3.1 Festucetalia valesiacae

5.3.1.1 Festucion valesiacae (incl. Festucion rupicolae)

- Astragalo-Festucetum rupicolae
- Cleistogeno-Festucetum rupicolae
- Cynodonto-Festucetum pseudovinae
- Cynodonto-Poetum angustifoliae
- Medicagini-Festucetum valesiacae
- Potentillo-Festucetum pseudodalmaticae
- Potentillo-Festucetum pseudovinae
- Pulsatillo-Festucetum rupicolae
- Salvio-Festucetum rupicolae

5.3.1.2 Cirsio-Brachypodion

- Lino tenuifolio-Brachypodietum pinnati
- Polygalo-Brachypodietum pinnati

5.3.1.3 Asplenio-Festucion pallentis

- Asplenio rutae-murariae-Melicetum ciliatae
- Asplenio septentrionali-Melicetum ciliatae
- lnulo-Festucetum pseudodalmaticae
- Minuartio-Festucetum pseudodalmaticae
- Poetum pannonicae

5.3.1.4 Bromo-Festucion pallentis

- Chrysopogono-Caricetum humilis
- Festuco pallenti-Brometum pannonici
- Poo badensi-Caricetum humilis

Seseli leucospermi-Festucetum pallentis

Seslerietum sadlerianae

Stipo-Festucetum pallentis

5.3.1.5 Saturejon montanae

Sedo sopianae-Festucetum dalmaticae

Serratulo radiatae-Brometum pannonici

5.3.1.6 Danthonio-Stipion tirsae

Campanulo macrostachyae-Stipetum tirsae

5.3.1.7 Artemisio-Kochion

Agropyro pectinati-Kochietum prostratae

Artemisietum ponticae-campestris

5.3.2 Brometalia erecti

5.3.2.1 Xerobromion

5.3.3 Festucetalia vaginatae

5.3.3.1 Festucion vaginatae

Festucetum vaginatae

Festuco vaginatae-Corynephoretum

5.3.3.2 Bromion tectorum

Brometum tectorum

5.4 Molinio-Arrhenatheretea

5.4.1 Molinietalia

5.4.1.1 Molinion

Junco-Molinietum

Molinio-Salicetum rosmarinifoliae

Succiso-Molinietum

5.4.1.2 Filipendulo-Petasition

Aconitetum gracilis

Aegopodio-Petasitetum

Chaerophyllo-Petasitetum

Equiseto-Chaerophylletum hirsuti

Filipendulo-Cirsietum oleracei

Filipendulo-Geranietum

Nasturtio-Petasitetum

5.4.1.3 Cnidion
5.4.1.4 Juncion acutiflori
5.4.1.5 Calthion
Aconitetum gracilis
5.4.1.6 Deschampsion caespitosae
Agrostio-Poetum trivialis
Agrostio-Typhoidetum
Carici-Alopecuretum pratensis
Cirsio cani-Festucetum pratensis
Deschampsietum caespitosae
5.4.2 Arrhenatheretalia
5.4.2.1 Arrhenatherion
Alopecuro-Arrhenatheretum
Pastinaco-Arrhenatheretum
Anthyllido-Festucetum pratensis
5.4.2.2 Polygono-Trisetion
Trisetetum flavescentis
5.4.2.3 Cynosurion
Festuco rubrae-Cynosuretum
Lolio-Cynosuretum
5.5 Festuco-Puccinellietea
5.5.1 Puccinellietalia
5.5.1.1 Puccinellion limosae
Bassietum sedoidis
Camphorosmetum annuae
Chenopodio-Puccinellietum limosae
Hordeetum hystricis
Pholiuro-Plantaginetum tenuiflorae
Puccinellietum limosae
5.5.1.2 Puccinellion peisonis
Lepidio crassifolii-Camphorosmetum
Lepidio crassifolii-Puccinellietum limosae
Lepidio crassifolii-Puccinellietum peisonis

Puccinellietum peisonis

5.5.1.3 Juncion gerardii

Agrostio-Caricetum distantis

Scorzonero parviflorae-Juncetum gerardii

Trifolio-Caricetum divisae

5.5.1.4 Beckmannion eruciformis

Agrostio-Alopecuretum geniculati

Agrostio-Alopecuretum pratensis

Agrostio-Beckmannietum

Agrostio-Glycerietum poiformis

5.5.2 Artemisio-Festucetalia

5.5.2.1 Festucion pseudovinae

Achilleo-Festucetum pseudovinae

Artemisio santonici-Festucetum pseudovinae

Lepidio crassifolii-Festucetum pseudovinae

Peucedano-Asteretum sedifolii

6. Forest edge heath and forb vegetation

6.1 Trifolio-Geranietea

6.1.1 Origanetalia vulgaris

6.1.1.1 Trifolion medii

6.1.1.2 Geranion sanguinei

6.2 Epilobietea angustifolii

6.2.1 Atropetalia (Epilobietalia angustifolii)

6.2.1.1 Epilobion angustifolii

Calamagrostietum epigei

Digitali-Calamagrostietum arundinaceae

Senecioni-Epilobietum angustifolii

6.2.1.2 Atropion

Atropetum belladonnae

Molinietum arundinaceae

7. Needleleaved forests and allied communities

7.1 Erico-Pinetea

- 7.1.1 Erico-Pinetalia
  - 7.1.1.1 Erico-Pinion
    - Chamaebuxo-Pinetum
- 7.2 Pulsatillo-Pinetea
  - 7.2.1 Pulsatillo-Pinetalia
    - 7.2.1.1 Cytiso-Pinion
      - Lino flavi-Pinetum
    - 7.2.1.2 Festuco vaginatae-Pinion
      - Festuco vaginatae-Pinetum sillvestris
- 7.3 Vaccinio-Piceetea
  - 7.3.1 Piceetalia
    - 7.3.1.1 Dicrano-Pinion
    - 7.3.1.2 Linnaeo-Piceion
      - 7.3.1.2.5 Vaccinio-Abietenion
        - Bazzanio-Abietetum

8. Broadleaved forests and woodlands
- 8.1 Salicetea purpureae
  - 8.1.1 Salicetalia purpureae
    - 8.1.1.1 Salicion elaeagni
      - Hippophae-Salicetum elaeagni
      - Myricario-Epilobietum dodonaei
    - 8.1.1.2 Salicion albae
      - Salicetum albae-fragilis
      - Salici-Populetum
    - 8.1.1.3 Salicion triandrae
      - Salicetum purpureae
      - Salicetum triandrae
- 8.2 Alnetea glutinosae
  - 8.2.1 Alnetalia glutinosae
    - 8.2.1.1 Alnion glutinosae
      - Dryopteridi-Alnetum
      - Fraxino pannonicae-Alnetum

Thelypteridi-Alnetum

8.3 Quercetea robori-petraeae

8.3.1 Quercetalia robori-petraeae

8.3.1.1 Quercion robori-petraeae (incl. Castaneo-Quercion)

Castaneo-Quercetum

Chrysanthemo-Quercetum

Genisto tinctoriae-Carpinetum

8.3.1.2 Pino-Quercion

Aulacomnio-Pinetum

Genisto nervatae-Pinetum

8.4 Querco-Fagetea

8.4.2 Quercetalia pubescentis-petraeae

8.4.2.1 Quercion pubescentis

8.4.2.2 Orno-Ostryon

Cotino-Quercetum pubescentis

Cotoneastro-Amelanchieretum

Fago-Ornetum

Inulo spiraeifolio-Quercetum pubescentis

Orno-Quercetum pubescentis

8.4.2.3 Aceri tatarico-Quercion

Aceri campestri-Quercetum

Aceri tatarico-Quercetum

Ceraso mahaleb-Quercetum pubescentis

Convallario-Quercetum

Corno-Quercetum

Dictamno-Tilietum cordatae

Festuco rupicolae-Ouercetum

Galatello-Quercetum

Junipero-Populetum albae

Poo pannonicae-Quercetum

Tilio-Fraxinetum excelsioris

Waldsteinio-Spiraeetum mediae

8.4.2.4 Quercion petraeae-cerris

Asphodelo-Quercetum roboris-cerris
Deschampsio-Quercetum petraeae-cerris
Genisto pilosae-Quercetum
Orno-Quercetum polycarpae
Potentillo micranthae-Quercetum
Quercetum petraeae-cerris

8.4.3 Fagetalia

8.4.3.1 Fagion sylvaticae

8.4.3.1.1 Luzulo-Fagion (incl. Deschampsio-Fagion)

Galio rotundifolio-Fagetum
Luzulo-Fagetum

8.4.3.1.2 Galio odoratae-Fagion

Aconito-Fagetum
Cyclamini-Fagetum
Laureolae-Fagetum
Melittio-Fagetum

8.4.3.1.3 Cephalanthero-Fagion

Seslerio hungaricae-Fagetum
Tilio-Sorbetum

8.4.3.1.4 Aceri-Fagion

Parietario-Aceretum
Phyllitidi-Aceretum

8.4.3.1.5 Tilio-Acerion

Mercuriali-Tilietum

8.4.3.1.6 Galio-Abietetum

Abieti-Fagetum

8.4.3.2 Carpinion betuli

Querco petraeae-Carpinetum
Querco robori-Carpinetum
Waldsteinio-Carpinetum

8.4.3.3 Alno-Ulmion

Aegopodio-Alnetum
Carici acutiformis-Alnetum

Carici brizoidis-Alnetum
Carici remotae-Fraxinetum
Fraxino pannonicae-Ulmetum
8.4.3.4 Aremonio-Fagion
8.4.3.4.1 Primulo-Fagenion
Asperulo taurinae-Carpinetum
Fraxino pannonicae-Carpinetum
Helleboro dumetorum-Carpinetum
Helleboro odori-Fagetum
Scutellario-Aceretum
Tilio tomentosae-Fraxinetum orni
Vicio oroboidis-Fagetum
8.4.3.4.2 Lamio orvalae-Fagenion
8.4.3.4.3 Lonicero-Fagenion
8.4.3.4.4 Ostryo-Fagenion
8.5 Franguletea
8.5.1 Pteridio-Rubetalia
8.5.1.1 Lonicero-Rubion silvatici
8.5.2 Salicetalia auritae
8.5.2.1 Salicion cinereae
Calamagrostio-Salicetum cinereae
Salici cinereae-Sphagnetum
Salici pentandrae-Betuletum
8.6 Rhamno-Prunetea
8.6.1 Prunetalia spinosae
8.6.1.1 Pruno-Rubion radulae
8.6.1.2 Pruno-Rubion ulmifolii
8.6.1.3 Carpino-Prunion
Coryletum avellanae
Pruno spinosae-Crataegetum
8.6.1.4 Berberidion
8.6.1.5 Prunion fruticosae
Amygdaletum nanae

Crataego-Cerasetum fruticosae

Phlomidi-Prunetum tenellae

8.6.2 Salicetalia arenariae

8.6.2.1 Salicion arenariae

8.6.3 Sambucetalia

8.6.3.1 Sambuco-Salicion capreae

Aegopodio-Sambucetum nigrae

Epilobio-Salicetum capreae

Fragario-Rubetum

Sambucetum racemosae

Solidagini-Cornetum sanguineae

Previously uncategorised species were categorised by Attila Lengyel.

Data source and citation:

Borhidi A. (1995) Social behaviour types, the naturalness and relative ecological indicator values of the higher plants in the Hungarian Flora. Acta Botanica Hungarica 39: 97-181.

Horváth, F., Dobolyi, K., Morschhauser, T., Lőkös, L., Karas, L., & Szerdahelyi, T. (1995) Flóra adatbázis 1.2. Taxon-lista és attribútum állomány. Vácrátót: MTA ÖBKI.

Sonkoly, J., Tóth, E., Balogh, N., Balogh, L., Bartha, D. ... Török, P. (2022) Introducing PADAPT 1.0, the Pannonian Database of Plant Traits. Journal of Vegetation Science (submitted manuscript)

Other references:

Bagi I. & Székely Á. (2006) Az *Elymus elongatus* (Host) Runemark, magas tarackbúza előfordulása a Kiskunság déli részén - a korábbi lelőhelyek rövid áttekintés. Botanikai Közlemények 93: 77–92.

Barina, Z., Somogyi, G. and Pifkó, D. (2020) Typification of names in the *Dianthus plumarius* group in the Carpatho-Pannonian region. Taxon 69: 161–169.

Bátori, Z., Erdős, L., & Somlyay, L. (2012) Euphorbia prostrata (Euphorbiaceae), a new alien in the Carpathian Basin. Acta Botanica Hungarica 54: 235–243.

Bauer, N. & Somlyay, L. (2015) A *Crepis mollis* (Jacq.) Asch. subsp. *hieracioides* (Waldst. & Kit.) Domin újrafelfedezése Magyarországon / Rediscovery of *Crepis mollis* (Jacq.) Asch. subsp. *hieracioides* (Waldst. & Kit.) Domin in Hungary. Kitaibelia 20: 150–156.

Borhidi, A. (1993) A magyar flóra szociális magatartás típusai, természetességi és relatív ökológiai értékszámai. JPTE Növénytani Tanszék, Pécs.

Czimber, Gy., Varga, Z. & Radics, L. (2008) Új mediterrán fajok a hazai gyomflórában: a fehér kányazsázsa (*Diplotaxis erucoides* (Torner) DC.) Növénytermelés 57: 253–265.

Csecserits, A. & Barabás, S. (2020) A labodalevelű szárnyaslibatop (*Cycloloma atriplicifolia*) újabb előfordulása a Kiskunság északi részén. Kitaibelia 25: 107–108.

Csiky, J. (2003) A *Cuscuta approximata* Babington Magyarországon (Cuscutaceae Dumort.). Kitaibelia 8: 75–80.

Csiky, J., Farkas, S., Király, G., Pál, R., Purger, D. & Tóth, I. Zs. (2005) A *Cirsium boujartii* (Pill. et Mitterp.) Schultz Bip. újrafelfedezése Magyarországon /Rediscovery of *Cirsium boujartii* (Pill. et Mitterp.) Schultz Bip. in Hungary/ Flora Pannonica 3: 69–78.

Csiky, J., Mesterházy, A., Szalontai, B. & Pótó-Oláh, E. (2010) A morphological study of *Ceratophyllum tanaiticum*, a species new to the flora of Hungary. Preslia 82: 247–259.

Dancza, I., Hoffmann, Z. P. & Doma, C. (2004) *Cyperus esculentus* (yellow nutsedge) - a new weed in Hungary. Zeitschrift für Pflanzenkrankheiten und Pflanzenschutz 19: 223–229.

Dítě, D., Eliáš, P. & Király, G. (2006) *Dactylorhiza lapponica* (Laest. ex Hartm.) Soó, a new taxon for Hungary. Flora Pannonica 4: 91–97.

Fekete, R., Mesterházy, A., Valkó, O. & Molnár V. A. (2018) A hitchhiker from the beach: the spread of the maritime halophyte *Cochlearia danica* along salted continental roads. Preslia 90: 23–37.

Hroudová, Z., Zákravský, P., Ducháček, M. & Marhold, K. (2007) Taxonomy, distribution and ecology of *Bolboschoenus* in Europe. Annales Botanici Fennici 44: 81–102.

Kerényi-Nagy, V. (2012) Piros áltermésű ritka galagonyafajok – *Crataegus* spp. in: Bartha D. (szerk.): Magyarország ritka fa- és cserjefajainak atlasza. Kossuth Kiadó, Budapest, pp. 185-193.

Király, G. & Király, A. (2004) Az *Agrimonia procera* Wallr. előfordulása Magyarországon. Flora Pannonica 2: 7–24.

Király, G. (2005) Kiegészítések a magyar adventív-flóra ismeretéhez II. Az *Epilobium ciliatum* Rafin. Magyarországon. Flora Pannonica 3: 27–39.

Király, G., Steták, D. & Bányász, Á. (2008) Spread of invasive macrophytes in Hungary. In: Rabitsch, W., F. Essl & F. Klingenstein (Eds.): Biological Invasions – from Ecology to Conservation. Neobiota 7: 123–130.

Király, G., Bidló, A., Takács, G., Eliáš, P., Melečková, Z. & Dítě, D. (2013) Remote locality of the littoral *Carex extensa* (Cyperaceae) in Hungary — long distance dispersal from coastal to inland salt marshes. Biologia 68: 872–878.

Király, G. & Király, A. (2018) Adatok és kiegészítések a magyar flóra ismeretéhez III. Botanikai Közlemények 105: 27–96.

Korda, M. (2013) Újabb adat a magyar adventívflóra ismeretéhez: az *Allium paradoxum* (M. Bieb.) G. Don 1827 Magyarországon. Kitaibelia 18: 31–34.

Király, G., Hohla, M., Süveges, K., Hábenczyus, A. A., Barina, Z., Király, A., Lukács, B. A., Türke, I. J. & Takács, A. (2019) Taxonomical and chorological notes 10 (98–110). Studia Botanica Hungarica 50: 391–407.

Kovács, D. & Mesterházy, A. (2015) A *Ceratochloa* (DC. et P. Beauv.) Hack. alnemzetség (*Bromus* L., Poaceae) hazai története és elterjedése. Kitaibelia 20: 44–47.

Kun, A. (2019) Az *Apium repens* császártöltési állományának monitorozása (2006–2015). Kitaibelia, 24: 1–8.

Mandák, B. & Prach, K. (2001) *Cycloloma atriplicifolia*, a new alien species in Hungary. Preslia 73: 153–160.

Mesterházy, A., Király, G. & Wallnöfer, B. (2011) On the occurrence of *Carex randalpina* B. Wallnöfer (Cyperaceae) in Hungary. Annalen des Naturhistorischen Museums in Wien. Serie B für Botanik und Zoologie 112: 177–180.

Mesterházy, A., Matus, G., Király, G., Szűcs, P., Török, P., Valkó, O., Pelles, G., Papp, V. G., Virók, V., Nemcsok, Z., Rigó, A., Hohla, M. & Barina, Z. (2017) Taxonomical and chorological notes 5 (59–68). Studia Botanica Hungarica 48: 263–275.

Molnár V. A. (szerk.) (2011) Magyarország orchideáinak atlasza. Kossuth Kiadó, Budapest, 504 pp.

Molnár, C. & Juhász, M. (2016) Az alacsony libatop (*Chenopodium pumilio* R.Br.) Zuglóban és új adatok Északkelet-Magyarország idegenhonos fajainak elterjedéséhez. Kitaibelia 21: 221–226.

Mosolygó-L, Á., Sramkó, G., Barabás, S., Czegledi, L., Jávor, A., Molnár, V. & Surányi, G. (2016) Molecular genetic evidence for allotetraploid hybrid speciation in the genus *Crocus* L. (Iridaceae). Phytotaxa 258: 121–136.

Pal, R.W. (2011) *Echinaria capitata* (Seslerieae, poaceae), a new grass species for the Hungarian flora. Acta Botanica Hungarica 53: 175–180.

Partosfalvi, P., Madarász, J. & Dancza, I. (2008) Az ázsiai gyapjúfű (*Eriochloa villosa* (Thunb.) Kunth) megjelenése Magyarországon. Növényvédelem 44: 297–304.

Pifkó, D. (2004) Adatok a hazai *Chamaecytisus* fajok ismeretéhez I. Flora Pannonica 2: 25–36.

Pinke, Gy., Czimber, Gy. & Pál, R. (1999) A *Chorispora tenella* (Pall.) DC. a Szigetközben. Kitaibelia 4: 287–288.

Pinke, Gy., Pál, R., Király, G., V., Szendrődi, V. & Mesterházy, A. (2006) The occurrence and habitat conditions of *Anthoxanthum puelii* Lecoq & Lamotte and other Atlantic-Mediterranean weed species in Hungary. Zeitschrift für Pflanzenkrankheiten und Pflanzenschutz Sonderheft 20: 587-596.

Pinke, G., Molnár, S., Garamvölgyi, V. & Barina, Z. (2012) The first occurrence of *Euphorbia davidii* in Hungary. (Új gyomnövény Magyarországon a Dávid-Kutyatej (*Euphorbia davidii* Subils).). Növényvédelem 48: 117–120.

Simon, T. (1992) A magyarországi edényes flóra határozója. Tankönyvkiadó, Budapest.

Simon, T. (2001) A havasi varázslófű (*Circaea alpina* L.) hazai cönológiája. Botanikai Közlemények 88: 107–116.

Simon, T. & Podani, J. (2007) Régi-új faj, az *Euphorbia segetalis* L. a magyar flórában. Kitaibelia 12: 121–123.

Solymosi, P. (2016) A magyarországi adventív flóra lappangó faja a sárgás varjúláb [*Coronopus didymus* (L.) Smith]. Növényvédelem 52: 598–599.

Somlyay, L. (2009) Occurrence of *Chamaesyce glyptosperma*, and a survey of the genus *Chamaesyce* (Euphorbiaceae) in Hungary. Annales historico-naturales Musei nationalis Hungarici 101: 23–32.

Soó, R. (1968) A magyar flóra és vegetáció rendszertani-növényföldrajzi kézikönyve, III. Akadémiai Kiadó, Budapest.

Soó, R. (1980) A magyar flóra és vegetáció rendszertani-növényföldrajzi kézikönyve, VI. Akadémiai Kiadó, Budapest.

Štech, M., Koutecký, P., Tribsch, A., Schratt-Ehrendorfer, L., Paszko, B. & Pachschwöll, C. (2020) *Calamagrostis purpurea* (Poaceae) – A long neglected boreal element, new for the flora of Austria. Neilreichia - Zeitschrift für Pflanzensystematik und Floristik Österreichs 11: 133–152.

Takács, A., Baráth, K., Csiky, J., Csikyné, R. É., Király, G., Nagy, T., Papp, V., Schmidt, D., Tamási, B. & Barina, Z. (2016) Taxonomical and chorological notes 3 (28–37). Studia Botanica Hungarica 47: 345–357.

Vidéki, R. (2005) *Cycloloma atriplicifolia* (Spreng.) J. M. Coulter és *Salsola collina* Pallas Magyarországon / *Cycloloma atriplicifolia* (Spreng.) J. M. Coulter und *Salsola collina* Pallas in Ungarn. / Flora Pannonica 3: 121–134.

Virók, V. & Farkas, R. (2008) Új növényfaj a hazai edényes flórában: a Haller-kövifoszlár (*Cardaminopsis halleri* (L.) Hayek). Kitaibelia 13: 29–33.

Vojtkó, A. (1996) Mirigyes fodorka (*Asplenium lepidum* C. Presl.) előfordulása a Bükk-hegységben. Kitaibelia 1: 25.

Wilhalm, T. (2009) *Digitaria ciliaris* in Europe. Willdenowia 39: 247–259.

Wirth, T. & Gyergyák, K. (2015) Az *Asparagus verticillatus* L. Magyarországon. Kitaibelia 20: 38–43.

Wolf, M. A. & Király, G. (2014) Euphorbia serpens (Euphorbiaceae), a new alien apecies in Hungary. Acta Botanica Hungarica 56: 243–250.

**Nativeness and invasion biology**

The dataset about nativeness and invasion biology was prepared by János Csiky[1], Lajos Balogh[2], István Dancza[3], Ferenc Gyulai[4], Gusztáv Jakab[5], Gergely Király B.[6], Éva Lehoczky[7], Attila Mesterházy[8], Patrícia Pósa[9] and Tamás Wirth[10].

[2]Savaria Múzeum, Természettudományi Osztály, 9700 Szombathely, Kisfaludy Sándor u. 9.,

[4]MATE Környezettudományi Doktori Iskola, 2100 Gödöllő, Páter Károly u. 1.,

[5]Magyar Agrár- és Élettudományi Egyetem Környezettudományi Intézet, Páter Károly utca 1., Gödöllő 2100; Bölcsészettudományi Kutatóközpont Régészeti Intézet, 1097 Budapest, Tóth Kálmán u. 4., B. épület 2. emelet,, ORCID id.: https://orcid.org/0000-0002-2569-5967

[7]MATE KRC Környezettudományi Intézet, Agroökológiai Csoport, 3200 Gyöngyös, Mátrai út 36., 3200 Gyöngyös, Mátrai út 36.; ATK Talajtani Intézet, 1022 Budapest, Herman Ottó út 15.,

[8]Ökológiai Kutatóközpont, Vizes Élőhelyek Funkcionális Ökológiai Kutatócsoport, 4026 Debrecen, Bem tér 18/C,

[9]Balaton-felvidéki Nemzeti Park Igazgatóság, Ökoturisztikai és Környezeti-nevelési Osztály, 8229 Csopak, Kossuth Lajos u. 16.; MATE Környezettudományi Doktori Iskola, 2100 Gödöllő, Páter Károly u. 1.,, ORCID id: https://orcid.org/0000-0003-3025-1313

The categories describing the main invasion biology attributes are partly consistent and/or comparable with attributes already available in databases in neighbouring countries.

**Nativeness**

Categories:

- Native species: Those species are considered native that have been present in Hungary throughout the Holocene (since ≈5000 BC) or were present in nature at least intermittently in subsequent periods (e.g., *Quercus robur*). Species that became extinct during the Holocene, before the Neolithic, and have later been re-introduced into Hungary with human assistance are not considered native.
- Alien species: Species that settled in Hungary after the appearance of major human population and the associated interventions (appearing after the beginning of the Neolithic period in the territory of Hungary, ≈ 5000 BC), mainly those that would have been unable to overcome the natural barriers that prevented their spread without human interventions (e.g., *Centaurea cyanus*).

- Cryptogenic species: Any species for which it is debatable whether it is native or alien in Hungary (e.g., *Abies alba*). Here we interpret this term only in terms of species' nativeness to Hungary.
- Not confirmed**:** Species with no confirming specimens from Hungary (e.g., *Pyrola media*).
- Planted only**:** Only planted, cultivated specimens or stands are known from Hungary (e.g., *Metasequoia glyptostroboides*).

**Residence time**

In case of alien species.

Categories:

- Archaeophytes: alien plant species for which there is historical, archaeological or archaeobotanical evidence of occurrence in Hungary before 1500 AD (e.g., *Agrostemma githago*).
- Neophytes: alien plant species that appeared in Hungary after 1500 AD (e.g., *Bromus catharticus*).
- Unclear: any plant species of doubtful indigenous origin (e.g., *Larix decidua*) or alien species which may have been established in Hungary before or after 1500 AD (e.g., *Ficus carica*).

  For any alien or cryptogenic species for which it is not possible to determine with certainty whether it is an archaeophyte or a neophyte, the following four possibilities apply:

  - native/alien?: probably native, less likely to be an alien species (and if it is alien, it is most likely to be archaeophyte) (e.g., *Spiraea crenata*).
  - alien/native?: probably alien species (most likely archaeophyte), less likely to be native (e.g., *Castanea sativa*).
  - archaeophyte/neophyte?: probably archaeophyte in Hungary, but this can only be stated with some uncertainty (it may be neophyte) (e.g., *Eranthis hyemalis*).
  - neophyte/archaeophyte?: probably neophyte in Hungary, but this can only be stated with some uncertainty (it may be archaeophyte) (e.g., *Chenopodium strictum*).

**Invasion status**

Following the traditional approach, we distinguish three main phases of establishment (casual, naturalised, and invasive species). Among invasive species we distinguish species that significantly change the species composition and structure of habitats (transformer species).

Categories:

- Casual species: species with casual (spontaneous or subspontaneous) occurrences (e.g., *Cucumis sativus*), excluding plants established by deliberate planting or seeding.
- Naturalised species: alien species with self-sustaining populations in Hungary (e.g., *Cabomba caroliniana*).
- Invasive species: alien species that reproduce rapidly and spread over large areas (e.g., *Erigeron annuus*).
- Transformer species: invasive species that significantly alter their biotic and/or abiotic environment, resulting in the permanent alteration or disappearance of the original vegetation (e.g., *Asclepias syriaca*).

**Introduction mode**

Categories:

- Accidental: plant species unintentionally introduced by humans, or species that have migrated from neighbouring countries on their own (e.g., *Plantago coronopus*).
- Deliberate (subspontaneous): alien species that got out from stands deliberately established in Hungary (e.g., *Opuntia humifusa*).
- Both: both cases (accidental and deliberate introductions) are known for the species (e.g., *Sorghum halepense*).
- Unknown: no reliable data is available for the species in this respect (e.g., *Ranunculus psilostachys*).

As a result of the classifications, the distribution of the attributes was as follows:

From the 2745 species included in PADAPT:

- 1839 taxa are native,
- 790 taxa are alien,
- 84 taxa are cryptogenic,
- 23 taxa have no confirmed occurrences in Hungary,
- 9 taxa occur only planted.

From the 874 alien and cryptogenic taxa:

- 301 taxa are archaeophytes,
- 447 taxa are neophytes,
- 126 taxa are unclear.

From the 126 unclear taxa:

- 67 taxa are native/alien?,
- 17 taxa are alien/native?,
- 24 taxa are archaeophyte/neophyte?,
- 18 taxa are neophyte/archaeophyte?

According to invasion status, from the 874 alien taxa:

- 318 taxa are casual,
- 471 taxa are naturalized,
- 63 taxa are invasive,
- 22 taxa are transformer.

According to the mode of introduction, from the 874 alien taxa:

- 442 taxa were introduced accidentally,
- 371 taxa were introduced deliberately,
- 38 taxa were introduced both ways,
- 23 taxa are unknown.

The dataset about nativeness and invasion biology was prepared by János Csiky, Lajos Balogh, Ferenc Gyulai, Gusztáv Jakab, Éva Lehoczky, Attila Mesterházy, Patrícia Pósa and Tamás Wirth.

Data source and citation:

Balogh, L., Dancza, I., Király, G. (2004) A magyarországi neofitonok időszerű jegyzéke, és besorolásuk inváziós szempontból. [Actual list of neophytes in Hungary and their classification according to their success.] In: Mihály, B., Botta-Dukát, Z. (eds.): Biológiai inváziók Magyarországon: Özönnövények. [Biological invasions in Hungary: Invasive plants.] – A KvVM Természetvédelmi Hivatalának tanulmánykötetei 9, TermészetBÚVÁR Alapítvány Kiadó, Budapest [in Hungarian]

Sonkoly, J., Tóth, E., Balogh, N., Balogh, L., Bartha, D. ... Török, P. (2022) Introducing PADAPT 1.0, the Pannonian Database of Plant Traits. Journal of Vegetation Science (submitted manuscript)

Other references:

Balogh, L., Dancza, I., Király, G. (mscr. 2016) A magyarországi flóra újjövevénynövényeinek jegyzéke [Catalogue of neophytes of Hungary], 2016. márc. 27. In: Balogh L., Dancza I., Gyulai F., Király G.: A magyarországi flóra jövevénynövényeinek jegyzéke. [Catalogue of alien plants of Hungary.] Kéziratos adatbázis (manuscript of database). Savaria Múzeum, Szombathely, 2016. márc. 27. [in Hungarian].

Balogh L., Gyulai F. (mscr. 2004) A magyarországi flóra ójövevénynövényei; előzetes jegyzék [Catalogue of arcaeophytes of Hungary; a preliminary list], 2004. júl. 15. In: Balogh L., Dancza I., Gyulai F., Király G.: A magyarországi flóra jövevénynövényeinek jegyzéke. [Catalogue of alien plants of Hungary.] Kéziratos adatbázis (manuscript of database). Savaria Múzeum, Szombathely, 2016. márc. 27. [in Hungarian].

Gyulai, F. (2010) Archaeobotany in Hungary. Seed, Fruit, Food and Beverages Remains in the Carpathian Basin: an Archaeobotanical Investigation of Plant Cultivation and Ecology from the Neolithic until the Late Middle Ages. Archaeolingua, Budapest

Pósa, P., Gyulai, F. (2019) A tájtörténet fontos forrásának, a Magyar Archaeobotanikai Adatbázisnak a bemutatása. In: Módosné Bugyi I. et al. (eds.): XII. tájtörténeti tudományos konferencia. Füleky György emlékkonferencia. Szarvas 2019. június 27-29. Érdi Rózsa Nyomda [in Hungarian]

Pósa, P., Vinogradov, Sz., Gyulai, F. (2020) The developement of weed vegetation in the Pannonian Basin as seen in the archaeobotanical records. Applied Ecology and Environmental Research 18: 7431-7444.

**Area type**

The area type of the species is based on the Flóra database (Horváth et al. 1995).

The area type categorization in the Flóra database was developed to be both simple and to provide a scientific basis for thorough analyses.

The area type categorization is as follows:

ADVENTIVE GROUP

**ADV******** Adventive elements

COSMOPOLITAN GROUP

**COS******** Cosmopolitan elements

EUROPEANGROUP

**CIR******** Circumpolar elements
**EUA******* Eurasian elements
**EUR****** European elements
**CEU***** Central-European elements

CONTINENTAL GROUP

**CON****** Continental elements
**PON***** Pontic elements
**PoM***** Pontic-Submediterranean elements
**PoP***** Pontic-Pannonic elements
**TUR***** Turanian elements

MEDITERRANEAN GROUP

**MED****** Mediterranean elements
**SME****** Submediterranean elements
**SMO***** Eastern-Submediterranean elements
**PaB***** Pannon-Balkan elements

**BAL*** Balkan elements
**ILL** Illyrian, Western-Balkan elements

ATLANTIC GROUP

**AsM**** (Sub)atlantic-Submediterranean elements
**SAT**** Subatlantic elements

NORDIC and MONTANE GROUP

**BOR**** Boreal, Nordic elements
**ALP**** Alpine elements
**CEA*** Central-European-Alpine elements
**ALB*** Alpine-Balkan elements
**CAR** Carpathian endemisms
**DAC** Eastern-Carpathean, Dacic elements

ENDEMIC GROUP

**PAN** Pannonian endemisms
**END* Local (super) endemisms

The number of asterisks (*) indicates the geographical size of the categories.

Data source and citation:

Horváth, F., Dobolyi, K., Morschhauser, T., Lőkös, L., Karas, L., & Szerdahelyi, T. (1995) Flóra adatbázis 1.2. Taxon-lista és attribútum állomány. Vácrátót: MTA ÖBKI. [Flora database 1.2, List of taxa and attributes.]

Sonkoly, J., Tóth, E., Balogh, N., Balogh, L., Bartha, D. ... Török, P. (2022) Introducing PADAPT 1.0, the Pannonian Database of Plant Traits. Journal of Vegetation Science (submitted manuscript)

### Conservation status and value (HUF), and year of protection

**Conservation status** laid down by the Decree 13/2001 KÖM currently in force (hereafter Decree) concerning plant and animal species protected or strictly protected in Hungary, strictly protected caves and plant and animal species of community interest in the European Union.

Categories:

- Not protected
- Protected (Annex I. and VII. to the Decree)
- Strictly protected (Annex I. to the Decree)

**Conservation value (HUF):** The conservation value of species laid down by the Decree, expressed in HUF.

**Year of protection** indicates the year of the decree or amending decree in which the species was first listed.

The dataset on conservation status and value was prepared by Kinga Bata.

Citation:

Sonkoly, J., Tóth, E., Balogh, N., Balogh, L., Bartha, D. ... Török, P. (2022) Introducing PADAPT 1.0, the Pannonian Database of Plant Traits. Journal of Vegetation Science (submitted manuscript)

### IUCN Red List and Hungarian Red List category

#### IUCN Red List category

The International Union for Conservation of Nature and Natural Resources (IUCN) is an international organisation for the conservation of natural values. Its main task is to encourage and promote the conservation of biodiversity and the sustainable use of nature's resources and the maintenance of ecosystems.

IUCN founded the Red List of Threatened Species in 1964, which has since become the most comprehensive global inventory of the conservation status of animal, plant, and fungus species.

Categories of the IUCN Red List:

- **EX** – Extinct
- **EW** – Extinct in the Wild
- **CR** – Critically Endangered
- **EN** – Endangered
- **VU** – Vulnerable
- **NT** – Near Threatened
- **LC** – Least Concern
- **DD** – Data Deficient
- **NE** – Not Evaluated

#### Hungarian Red List category

Categorisation of the species based on Király (2007).

Categories and their description:

- **EX** – Extinct. A taxon is extinct if its last individual is definitely dead (all its previous habitats have been physically destroyed); or no specimens have been detected in the last 50 years during thorough and systematic research covering all its known habitats and additional presumed habitats.

**EW** – Extinct in the wild. A taxon is extinct in the wild if it occurs exclusively in cultivation or it only has introduced population(s) far from its original range; and no specimens have been detected in the last 50 years during thorough and systematic research covering all its known habitats and additional presumed habitats.

**CR** – Critically endangered. A taxon is critically endangered if it is particularly threatened by the risk of extinction in its natural environment.

**EN** – Endangered. A taxon is endangered if it is most likely to be threatened by the risk of extinction in its natural environment.

**VU** – Vulnerable. A taxon is vulnerable if it is threatened by a lower risk of extinction in its natural environment.

**NT** – Near threatened. A taxon is near threatened if it does not currently fall into any of the above categories, but, based on the current situation, it is likely to meet at least the criteria for the vulnerable category in the near future.

**DD** – Data deficient. A taxon is data deficient if no direct or indirect data are available on the current distribution or status of their populations, therefore the degree of vulnerability cannot be determined. Taxa that are not sufficiently studied in Hungary but certainly do not fall into the above vulnerability categories were not categorised as data deficient. In other words, taxa considered as data deficient are probably threatened by extinction, but the degree of vulnerability cannot be determined due to taxonomic uncertainties or to the lack of data on their localities.

The dataset on Red List categories was prepared by Kinga Bata.

Data source and citation:

Király G. (ed.) (2007) Vörös Lista. A magyarországi edényes flóra veszélyeztetett fajai. [Red list of the vascular flora of Hungary], Sopron

Sonkoly, J., Tóth, E., Balogh, N., Balogh, L., Bartha, D. ... Török, P. (2022) Introducing PADAPT 1.0, the Pannonian Database of Plant Traits. Journal of Vegetation Science (submitted manuscript)

**Simon-type conservation value category (TVK)**

Simon-type conservation value category of the species based on the Flóra database (Horváth et al. 1995).

Categories of the evaluation system:

Group I – species indicating naturalness

**U** – unique and very rare species
**KV** – species strictly protected in Hungary
**V** – species protected in Hungary
**E** – native edificator species (dominant species in plant communities)
**K** – native accessorial species
**TP** – natural pioneer species

Group II – species indicating degradation

**TZ** – disturbance-tolerant native species
**A** – adventive species
**G** – cultivated species
**GY** – weeds

Data source and citation:

Simon T. (1995) A hazai edényes flora természetvédelmi-érték besorolása. Abstracta Botanica 12: 1-23.

Horváth, F., Dobolyi, K., Morschhauser, T., Lőkös, L., Karas, L., & Szerdahelyi, T. (1995) Flóra adatbázis 1.2. Taxon-lista és attribútum állomány. Vácrátót: MTA ÖBKI. [Flora database 1.2, List of taxa and attributes.]

Sonkoly, J., Tóth, E., Balogh, N., Balogh, L., Bartha, D. ... Török, P. (2022) Introducing PADAPT 1.0, the Pannonian Database of Plant Traits. Journal of Vegetation Science (submitted manuscript)
