## Appendix S5 for "PADAPT 1.0 – the Pannonian Database of Plant Traits"

**Supporting Information to the paper Sonkoly, J. et al. Introducing PADAPT 1.0, the Pannonian Database of Plant Traits. *Journal of Vegetation Science.***

**Appendix S5. Descriptions and data sources of the attributes in the 'Ecological indicator values' group.**

### Ellenberg's ecological indicator values

#### Ellenberg's L: Light indicator value (Lichtzahl)

1 – Plants of deep shade
2 – Species intermediate between 1 and 3
3 – Shade plants, mostly <5% relative illumination
4 – Species intermediate between 3 and 5
5 – Semi-shade plants, >10% relative illumination, but seldom in full light
6 – Species intermediate between 5 and 7
7 – Plants of well-lit habitats tolerating partial shade
8 – High-light plants, seldom occurring at <40% relative illumination
9 – Plants of full light, occurring mostly in full sun

#### Ellenberg's T: Temperature indicator value (Temperaturzahl)

1 – Species indicating a cold climate, only in high mountain areas, in the alpine and nival regions
2 – Species intermediate between 1 and 3
3 – Species indicating a cool climate, mainly in subalpine regions
4 – Species intermediate between 3 and 5, especially montane and high montane species
5 – Species indicating a moderately warm climate, from lowlands to montane regions, with an emphasis on submontane regions
6 – Species intermediate between 5 and 7 (from planar to collin regions)
7 – Species indicating a cold climate, in northern Central Europe only in relatively warm lowlands
8 – Species intermediate between 7 and 9, with an emphasis on submediterranean areas
9 – Species indicating an extremely warm climate, from the Mediterranean area to warmest places in southern Central Europe

#### Ellenberg's K: Continentality indicator value (Kontinentalitätszahl)

1 – Euoceanic species, in Central Europe only with a few outposts
2 – Oceanic species, with an emphasis on Western Europe, including western Central Europe
3 – Species intermediate between 2 and 4, i.e., in large parts of Central Europe
4 – Suboceanic species, centered in Central Europe, spreading eastward
5 – Intermediate species, slightly suboceanic to slightly subcontinental
6 – Subcontinental species, with an emphasis on eastern Central Europe and Eastern Europe
7 – Species intermediate between 6 and 8
8 – Continental species, only at special locations spanning from Eastern to Central Europe
9 – Eucontinental species, absent in western Central Europe and rare in Eastern Europe

**Ellenberg's F: Soil moisture indicator value (Feuchtezahl) (1–12)**

1 – Species of extremely dry habitats, on soils that often dry out completely
2 – Species intermediate between 1 and 3
3 – Species indicating dry habitats, occurring mostly in dry soils
4 – Species intermediate between 3 and 5
5 – Species indicating soil moisture, mostly on soils with average water content
6 – Species intermediate between 5 and 7
7 – Species of constantly moist but not wet soils
8 – Species intermediate between 7 and 9
9 – Species indicating high soil moisture, mostly occurring on water-saturated soils
10 – Species of shallow-water habitats that may lack standing water in some periods of the year
11 – Species that root underwater and are either floating on the water surface or partially emerge above the surface
12 – Submerged plants that permanently live underwater

**Ellenberg's R: Soil reaction indicator value (Reaktionszahl) (1–9)**

1 – Species indicating extreme acidity, always found on acidic soils
2 – Species intermediate between 1 and 3
3 – Species indicating acidity but sometimes occurring on nearly neutral soil
4 – Species intermediate between 3 and 5
5 – Species indicating slightly acidic soils, rarely occurring in highly acidic soils
6 – Intermediate species between 5 and 7
7 – Species indicating slightly basic soils, never occurring in highly acidic habitats
8 – Intermediate species between 7 and 9
9 – Species indicating basic conditions, always found on calcareous and other basic soils

**Ellenberg's N: Nutrient availability indicator value (Nährstoffzahl) (1–9)**

1 – Species indicating soil with extremely low nutrient availability
2 – Species intermediate between 1 and 3
3 – Species of moderately infertile soils
4 – Species intermediate between 3 and 5
5 – Species indicating intermediate soil fertility
6 – Species intermediate between 5 and 7
7 – Species of habitats with mostly high fertility
8 – Species intermediate between 7 and 9
9 – Species indicating extremely rich soils

**Ellenberg's S: Salt content indicator value (Salzzahl) (0–9)**

0 – Species absent from saline soils
1 – Slightly salt-tolerant species able to persist in the presence of salt
2 – Species occurring in both saline and non-saline soils
3 – Species of mostly saline, coastal habitats that also occur on non-saline soils
4 – Species occurring in habitats with moderately saline soils, salt meadows and upper salt marshes

5 – Species of saline habitats, such as cliffs receiving salt spray
6 – Species of mostly highly saline habitats, salt marshes
7 – Species of highly saline soils of lower salt marshes
8 – Species of soils mostly inundated by sea water
9 – Species of extremely saline conditions, where sea water evaporates and precipitates salt

Data source and citation:

Ellenberg, H., Weber, H. E., Düll, R., Wirth, V., Werner, W. & Paulissen, D. (1991) Zeigerwerte von pflanzen in Mitteleuropa. Scripta Geobotanica 18.

Sonkoly, J., Tóth, E., Balogh, N., Balogh, L., Bartha, D. ... Török, P. (2022) Intorducing PADAPT 1.0, the Pannonian Database of Plant Traits. Journal of Vegetation Science (submitted manuscript)

### Zólyomi's ecological indicator values

#### Zólyomi's T: temperature requirement (1-7, 0)

1 – in accordance with the alpine, or tundra belt
2 – in accordance with the boreal tundra belt
3 – in accordance with the taiga belt
4 – in accordance with the mixed forest belt
5 – in accordance with the broad-leaved forest belt
6 – in accordance with the submediterranean woodland belt
7 – in accordance with the mediterranean, atlantic evergreen belt
0 – indifferent

Atlantic and continental climatic requirements of species are denoted by 'a' or 'k' after the TZ value.

#### Zólyomi's W: moisture requirement (0-11)

0 – in accordance with extremely dry habitats
1 – in accordance with habitats with long dry period
2 – in accordance with dry habitats
3 – in accordance with semi-dry habitats
4 – in accordance with semi-humid habitats
5 – in accordance with fresh habitats
6 – in accordance with moderately moist soils
7 – in accordance with moist-wet soils
8 – in accordance with wet soils
9 – in accordance with inundated soils
10 – in accordance with wetland, floating vegetation
11 – in accordance with water bodies, water plants

#### Zólyomi's R: soil requirement, soil reaction (1-5, 0)

1 – in accordance with acidic soils, calciphobe plants

2 – in accordance with moderately acidic soils
3 – in accordance with neutral soils
4 – in accordance with slightly basic soils
5 – in accordance with basic soils, basiphilous plants
0 – indifferent

Data source and citation:

Zólyomi, B., Baráth, Z., Fekete, G., Jakucs, P., Kárpáti, I., Kárpáti, V., ... & Máthé, I. (1967) Einreihung von 1400 Arten der ungarischen Flora in ökologische Gruppen nach TWR-Zahlen. Fragm. Bot. Mus. Hist. Nat. Hung, 4, 101-142.

Horváth, F., Dobolyi, K., Morschhauser, T., Lőkös, L., Karas, L., & Szerdahelyi, T. (1995) Flóra adatbázis 1.2. Taxon-lista és attribútum állomány. Vácrátót: MTA ÖBKI. [Flora database 1.2, List of taxa and attributes.]

Sonkoly, J., Tóth, E., Balogh, N., Balogh, L., Bartha, D. ... Török, P. (2022) Introducing PADAPT 1.0, the Pannonian Database of Plant Traits. Journal of Vegetation Science (submitted manuscript)

**Soó's ecological indicator values**

**Soó's T: temperature requirement (1–5, 0)**

1 – highly cold-tolerant plants, arctic, alpine species
2 – cold-tolerant plants
3 – slightly cold-tolerant plants
4 – cold-sensitive, heat-demanding plants
5 – highly heat-demanding plants
0 – indifferent species

**Soó's F: soil moisture requirement (1-5, 0)**

1 – plants living in extremely dry habitats
2 – plants mostly living in dry habitats. but occasionally occurring in mesophilous habitats
3 – plants living in moderately wet habitats
4 – plants living in wet habitats that sometimes dry out
5 – plants living in constantly wet habitats
0 – indifferent species

**Soó's R: soil reaction, or Calcium-demand (1-5, 0)**

1 – calciphobe plants living in highly acidic soils
2 – plants living in moderately acidic soils, avoiding Ca in the soil
3 – plants living in neutral soils, avoiding Ca in the soil
4 – plants living in neutral soils
5 – plants living in basic soils

0 – indifferent species

**Soó's N: nitrogen requirement (1-5, 0)**

1 – plants always living in nitrogen-poor habitats
2 – plants mostly living in nitrogen-poor habitats
3 – plants requiring average nitrogen amounts
4 – plants living in nitrogen-rich, well fertilized soils
5 – plants living in highly nitrogen-rich, overfertilized soils
0 – indifferent species

Data source and citation:

Soó, R. (1964-1980) A magyar flóra és vegetáció rendszertani-növényföldrajzi kézikönyve, I-VI. Akadémiai Kiadó, Budapest.

Horváth, F., Dobolyi, K., Morschhauser, T., Lőkös, L., Karas, L., & Szerdahelyi, T. (1995) Flóra adatbázis 1.2. Taxon-lista és attribútum állomány. Vácrátót: MTA ÖBKI. [Flora database 1.2, List of taxa and attributes.]

Sonkoly, J., Tóth, E., Balogh, N., Balogh, L., Bartha, D. ... Török, P. (2022) Intorducing PADAPT 1.0, the Pannonian Database of Plant Traits. Journal of Vegetation Science (submitted manuscript)

**Borhidi's ecological indicator values**

In case of eurytopic species Borhidi's scale typically uses the middle value of 5, in contrast with Ellenberg's approach, which denotes indifferent species with 0 or X.

**Borhidi's T: heat supply of the habitat where the species occurs (0–8)**

0 – in accordance with the subnival or supraboreal belt
1 – in accordance with the alpine, boreal or tundra belt
2 – in accordance with the subalpine or subboreal belt
3 – in accordance with montane needle-leaved forest or taiga belt
4 – in accordance with the mesophilous broad-leaved forest belt
5 – in accordance with the submontane broad-leaved forest belt
6 – in accordance with the thermophilous forest or forest-steppe belt
7 – in accordance with the submediterranean woodland and grassland belt
8 – in accordance with the eumediterranean evergreen belt

**Borhidi's W: occurrence in relation to soil moisture or the water table (1–12)**

1 – plants of extremely dry habitats or bare rock surfaces
2 – xero-indicators of habitats with a long dry period
3 – xero-tolerant plants occasionally occurring on wet soils
4 – plants of semi-dry habitats
5 – plants of semi-humid habitats of mesic conditions

6 – plants of usually wet soils
7 – plants of moist but well-aerated soils
8 – plants of moist soils tolerating short flooding
9 – plants of wet, poorly aerated soils
10 – plants of frequently flooded soils
12 – water plants with floating or partly emergent leaves
12 – aquatic plants, usually fully submersed in water

**Borhidi's R: soil reaction of the habitat**

1 – extremely acidophilous, explicitly calciphobe plants
2 – intermediate type between RB 1 and RB 3
3 – acidofrequent plants, mostly on acidic soils
4 – moderately acidophilous plants
5 – plants of slightly acidic soils
6 – plants mostly on neutral soils, sometimes also on acidic and basic soils, generally widely tolerant, more or less indifferent plants
7 – basifrequent plants, mostly on basic soils
8 – basiphilous plants
9 – explicitly calciphilous plants and ultrabasic specialists

**Borhidi's N: in relation to the ammonia and nitrate supply of the habitats (1–9)**

1 – only in soils extremely poor in mineral nitrogen
2 – plants of habitats very poor in nitrogen
3 – plants of moderately oligotrophic habitats
4 – plants of submesotrophic habitats
5 – plants of mesotrophic habitats
6 – plants of moderately nutrient rich habitats
7 – plants of soils rich in mineral nitrogen
8 – N-indicator plants of fertilized soils
9 – plants only on hyperfertilized soils, extremely rich in nitrogen

**Borhidi's L: in relation to relative light intensity during the summer**

1 – full-shade plants, often receiving less than 1% of the full light
2 – very shade-tolerant plants
3 – shade plants that also occur in habitats receiving more light
4 – shade – semi-shade plants
5 – semi-shade plants
6 – semi-shade – semi-light plants
7 – semi-light plants mostly living in full light, but also somewhat shade-tolerant
8 – high-light plants
9 – full-light plants of open habitats

**Borhidi's C: in relation to the distribution of plants according to degree of continentality of the climate (1–9)**

1 – euoceanic species, reaching Central Europe (CE) only in the West, not reaching Hungary

2 – oceanic species, mainly in Western Europe and in the western part of CE
3 – oceanic-suboceanic species with a center of distribution in CE
4 – suboceanic species, mainly in CE but expanding to the East
5 – intermediate species with slight suboceanic-subcontinental character
6 – subcontinental species with a center of distribution in eastern CE
7 – continental-subcontinental species with a center of distribution in Eastern Europe
8 – continental species reaching only the eastern part of CE
9 – eucontinental species with a center of distribution in Siberia and Eastern Europe

**Borhidi's S: in relation to the salt concentration of the soils (0–9)**

0 – species not occurring in salty or alkaline soils
1 – salt-tolerant plants, mainly in non-saline soils [<0.1% $Cl^-$]
2 – oligohaline plants, in soils with extremely low chloride content [0.05-0.3% $Cl^-$]
3 – beta-mesohaline plants, in soils with low chloride content [0.3-0.5% $Cl^-$]
4 – alfa/beta-mesohaline plants, in soils with intermediate chloride content [0.5-0.7% $Cl^-$]
5 – alfa-mesohaline plants, in soils with intermediate chloride content [0.7-0.9% $Cl^-$]
6 – alfa-mesohaline to polyhaline plants, in soils with intermediate to high chloride content [0.9-1.2% $Cl^-$]
7 – polyhaline plants, in soils with high chloride content [1.2-1.6% $Cl^-$]
8 – euhaline plants, in soils of very high chloride content [1.6-2.3% $Cl^-$]
9 – euhaline to hypersaline plants, in soils with extremely high chloride content [>2.3% $Cl^-$]

Data source and citation:

Borhidi A. (1995) Social behaviour types, the naturalness and relative ecological indicator values of the higher plants in the Hungarian Flora. Acta Botanica Hungarica 39: 97-181.

Sonkoly, J., Tóth, E., Balogh, N., Balogh, L., Bartha, D. ... Török, P. (2022) Intorducing PADAPT 1.0, the Pannonian Database of Plant Traits. Journal of Vegetation Science (submitted manuscript)
