## Appendix S6 for "PADAPT 1.0 – the Pannonian Database of Plant Traits"

**Supporting Information to the paper Sonkoly, J. et al. Introducing PADAPT 1.0, the Pannonian Database of Plant Traits. *Journal of Vegetation Science.***

**Appendix S6. Descriptions and data sources of the attributes in the 'Leaf traits' group.**

**Leaf Area (LA)**

The surface area of one side of freshly collected intact and mature leaves. Unit of measurement: $mm^2$.

PADAPT includes leaf trait data from Hungary, based on the data of Lhotsky et al. (2016), E-Vojtkó et al. (2020), McIntosh-Buday et al. (2022) and Gyalus et al. (2022).

**Leaf Dry Matter Content (LDMC)**

Leaf dry matter content is a measure of tissue density, given as the ratio of leaf dry mass to fresh mass expressed in mg/g.

PADAPT includes leaf trait data from Hungary, based on the data of Lhotsky et al. (2016), E-Vojtkó et al. (2020), McIntosh-Buday et al. (2022) and Gyalus et al. (2022).

**Specific Leaf Area (SLA)**

Specific leaf area is the ratio of leaf area to leaf dry weight. SLA = leaf area / leaf dry weight expressed in $mm^2/mg$.

PADAPT includes leaf trait data from Hungary, based on the data of Lhotsky et al. (2016), E-Vojtkó et al. (2020), McIntosh-Buday et al. (2022) and Gyalus et al. (2022).

**Leaf fresh mass**

The fresh mass of freshly collected mature and intact leaves expressed in grams (g).

PADAPT includes leaf trait data from Hungary, based on the data of Lhotsky et al. (2016), E-Vojtkó et al. (2020), McIntosh-Buday et al. (2022) and Gyalus et al. (2022).

**Leaf dry mass**

The mass of intact mature leaves dried to weight constancy, expressed in milligrams (mg).

PADAPT includes leaf trait data from Hungary, based on the data of E-Vojtkó et al. (2020) and McIntosh-Buday et al. (2022).

Data source and citation:

The range of sources that is to be cited is up to the user based on the range of the data used.

E-Vojtkó, A., Balogh, N., Deák, B., Kelemen, A., Kis, Sz., Kiss, R., Lovas-Kiss, Á., Löki, V., Lukács, K., Molnár, V. A., Nagy, T., Sonkoly, J., Süveges, K., Takács, A., Tóth, E., Tóth, K., Tóthmérész, B., Török, P., Valkó, O., Vojtkó, A., & Lukács, B. A. (2020) Leaf trait records of vascular plant species in the Pannonian flora with special focus on endemics and rarities. Folia Geobotanica 55: 73–79.

Gyalus, A., Barabás, S., Berki, B., Botta-Dukát, Z., Kabai, M., Lengyel, A., Lhotsky, B., Csecserits, A. (2022) Plant trait records of the Hungarian and Serbian flora and methodological description of some hardly measurable plant species. Acta Botanica Hungarica 64: 451-454.

Lhotsky, B., Csecserits, A., Kovács, B. & Botta-Dukát, Z. (2016) New plant trait records of the Hungarian flora. Acta Botanica Hungarica 58: 397–400.

McIntosh-Buday, A., Sonkoly, J., Takács, A., Balogh, N., Kovacsics-Vári, G., Teleki, B., Süveges, K., Tóth, K., Hábenczyus, A. A., Lukács, B. A., Lovas-Kiss, Á., Löki, V., Tomasovszky, A., Tóthmérész, B., Török, P., Tóth, E. (2022) New data of plant leaf traits from Central Europe. Data in Brief 108286

Sonkoly, J., Tóth, E., Balogh, N., Balogh, L., Bartha, D. ... Török, P. (2022) Introducing PADAPT 1.0, the Pannonian Database of Plant Traits. Journal of Vegetation Science (submitted manuscript)
